# Programmable pathway profiles reveal signaling principles of TGF-β superfamily receptors

**DOI:** 10.1101/2025.06.15.659618

**Authors:** Bo Gu, James M. Linton, Brice Graham Hendrickson, Hengyu Li, Ron Hadas, Gal Manella, Jan Gregrowicz, Bryan Anggito, Rong Lu, Michael B. Elowitz

## Abstract

The Transforming Growth Factor beta (TGF-β) superfamily, like other biological pathways, relies on families of co-expressed, partially redundant protein components, such as receptor subunits. The inability to systematically modulate multi-gene component expression profiles has made it difficult to understand how components and sets of components collectively process information. To overcome this, we developed *Pathway Sculptor*, a dCas12a-based epigenetic editing system that achieves simultaneous same-cell knockdown of at least twelve target genes. Programming TGF-β receptor profiles, by knocking down different receptor subsets, revealed functional interactions between the canonical BMP and TGF-β pathway branches. Unexpectedly, signaling within each branch depended on receptors in the opposite branch. Further, different receptor subsets played distinct roles: ACVR-class receptors modulated signaling magnitude, whereas BMPRs and TGFBRs discriminated among ligand variants. These results show how the two branches of the TGF-β superfamily collaboratively process signals, and establish *Pathway Sculptor* as a general platform for high-order combinatorial perturbation.

## Introduction

Cell-cell communication pathways, such as Transforming Growth Factor beta (TGF-β), Wnt, and Receptor Tyrosine Kinase (RTK) signaling systems, comprise families of ligand, receptor, and transducer variants. Different cell types express distinct but overlapping combinations of these components^1^, many of which exhibit partial redundancy or cross-regulatory interactions. This makes it difficult to determine the role of individual components, how specific sets of components combine to generate complex functions, and how information flows through pathway components in specific cellular contexts (**Fig. 1A**).

**Fig. 1.**
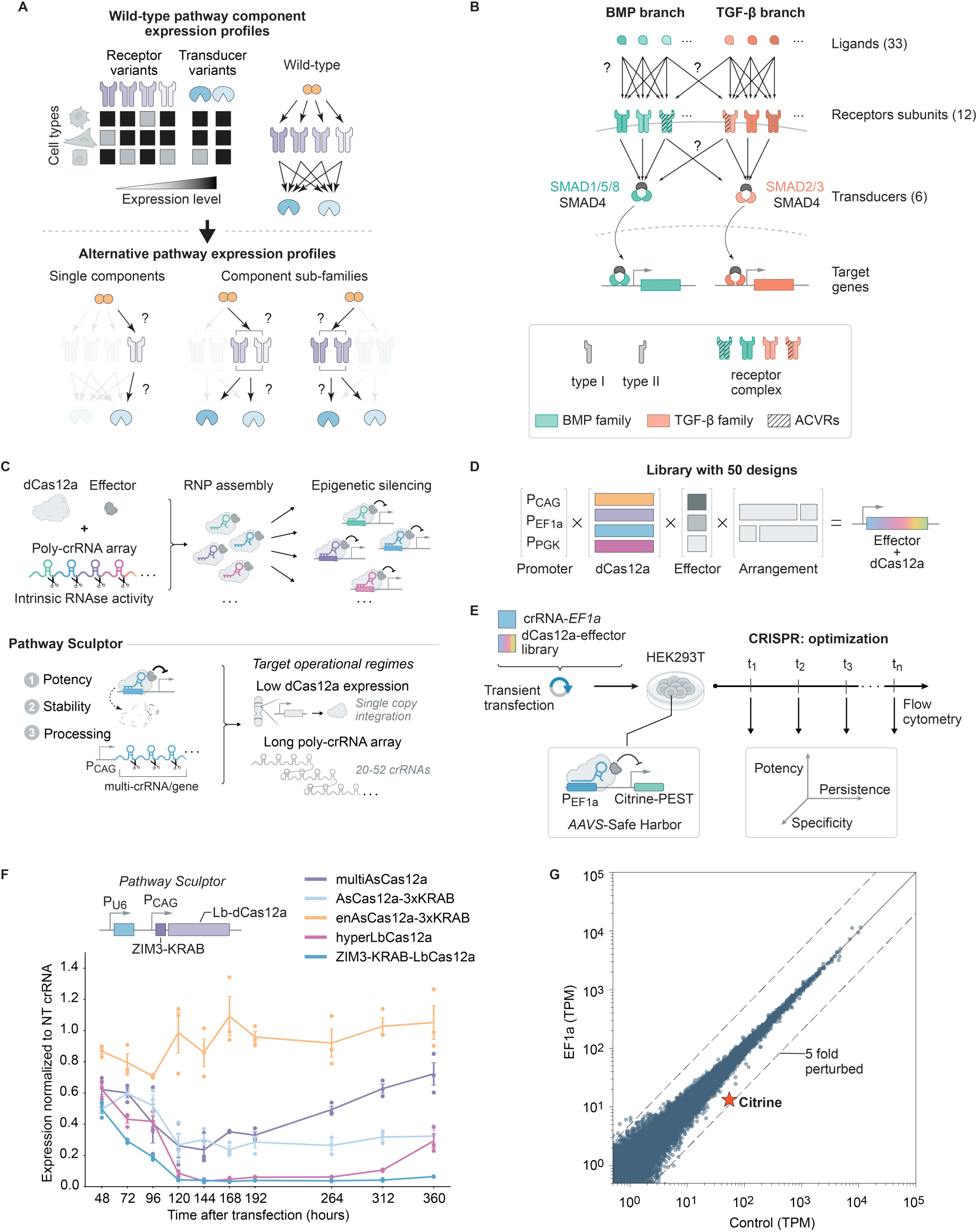
Identification of an enhanced dCas12a-effector variant. **(A)** Cell-cell communication pathways operate through families of component variants that are co-expressed in individual cell types (schematic). **(B)** Schematic of the overall TGF-β signaling system. BMP-family ligands (green) and TGF-β-family ligands (red) signal through distinct combinations of type I and type II receptors and subsequently activate the SMAD1/5/8–SMAD4 and SMAD2/3–SMAD4 transducers, respectively. Activated SMAD transducers translocate into the nucleus to regulate gene expression. Each receptor subunit is simultaneously associated with two orthogonal categorical labels: its conventional assignment to the BMP (green) or TGF-β (red) receptor family, and whether it belongs to the ACVR class (hatched) or not. Note that these receptor subunits are illustrative; a more specific depiction is introduced in Figure 3. **(C)** Top: Schematic of the dCas12a-based CRISPRi system. Bottom: Joint engineering of dCas12a–effector potency, component stability, and poly-crRNA array architecture constitutes *Pathway Sculptor*, which expands the performance space of dCas12a-based CRISPRi to enable large-scale multi-gene repression at restricted expression levels. **(D)** Design of dCas12a-effector variants library composed of combinations of promoter variants, dCas12a variants, transcriptional effectors, and construct arrangement variants. **(E)** Experimental workflow for systematic dCas12a-effector library screening and multi-dimensional optimization. **(F)** Benchmark of enhanced dCas12a-effector variant against existing dCas12a-based CRISPRi platforms. Each data point denotes the median normalized GFP intensity of an independent experiment. Error bars denote the mean ± s.e.m. across three independent biological replicates. **(G)** RNA-seq read-counts comparing conditions transfected with non-targeting (NT) crRNA vs EF1α-targeting crRNA. *n* = 3 biologically independent RNA-seq experiments. Dots represent transcripts; star denotes citrine transcript.

The TGF-β superfamily pathway exemplifies these challenges (**Fig. 1B**). In mammals, it includes over thirty ligands that signal through heterotetrameric receptor complexes assembled from two type I (of seven) and two type II (of five) receptor subunits. Once assembled, receptors can phosphorylate downstream SMAD transcription factors to regulate gene expression^2^. Historically, components of the pathway have been partitioned into two branches — BMP, which signals through SMAD1/5/8, and TGF-β, which signals through SMAD2/3. However, the two branches share an overlapping set of pathway components including the ACVR receptors and SMAD4, which could allow inter-branch crosstalk.

Consistent with this possibility, the two branches exhibit context-dependent interactions across many processes. For example, Nodal (TGF-β branch) and BMP4 (BMP branch) antagonize each other in establishing mesoderm^3^, while BMP2 and activin synergize to induce follicle-stimulating hormone synthesis in pituitary gonadotropes^4^. However, whether such interactions reflect an intrinsic property of the pathway architecture remains unclear. More specifically, it remains unclear to what extent ligand signaling is restricted to its respective branches, and how individual receptor pairs and specific receptor subsets mediate ligand signaling. Resolving these issues would help explain why the same ligand produces different outputs across cellular contexts, why receptor expression profiles vary across tissues, and why disease perturbations of the pathway are difficult to predict.

Addressing these questions requires the ability to remove or modulate specific combinations of receptors within the same cell. CRISPR-based tools offer a promising route towards this goal by allowing programmable genetic perturbation: across multiple platforms, single and pairwise gene activation and repression have become routine^5–7^, with higher-order perturbations demonstrated in select contexts^8–10^. However, existing approaches have not achieved reliable and potent same-cell multiplexed gene perturbation beyond two or three targets, which is insufficient to systematically dissect information flow in many major pathways.

Here, we introduce *Pathway Sculptor*, a scalable dCas12a-based epigenetic editing system that can achieve large-scale multi-gene perturbation in single cells, and how it can be used to dissect information flow within the TGF-β superfamily pathway. The engineering involved jointly optimizing the epigenetic effectors, component stability, and crRNA array architecture, with the combination of all three required for reliable higher-order performance. The resulting system, delivered as a single genomically integrated copy, achieved simultaneous knockdown of at least twelve targets, with ≥70% transcript reduction per target. Using *Pathway Sculptor*, we systematically programmed various TGF-β receptor profiles and analyzed the resulting signaling responses. The results revealed an unexpected reciprocal dependence of each branch on the other’s receptors and functional specialization of receptor subsets in controlling signaling magnitude and in ligand identity discrimination. These findings reveal how varying receptor expression profiles can control signaling magnitude, inter-branch crosstalk, and ligand discrimination. More generally, these results demonstrate how programming multi-component expression profiles using *Pathway Sculptor* can provide fundamental insights into central biological pathways.

## Results

### Identification of a dCas12a-effector variant with enhanced potency

The ability to simultaneously perturb twelve receptors at the single-cell level would enable systematic analysis of signaling capabilities of arbitrary TGF-β receptor profiles. CRISPR-Cas12a is particularly well suited for same-cell, multi-gene perturbation: its intrinsic RNase activity allows it to process poly-crRNA arrays into individual crRNAs without accessory proteins, and its compact 19-nt direct repeat enables efficient design of large multi-targeting arrays^11^. Its DNase-dead form, dCas12a, can be fused with epigenetic silencing domains to repress target genes without inducing genotoxic double-stranded DNA breaks, an approach collectively termed dCas12a-based CRISPRi^12–14^ (**Fig. 1C**, top).

Targeting twelve receptors simultaneously with dCas12a-based CRISPRi requires the use of long poly-crRNA arrays. Furthermore, high dCas12a expression can be cytotoxic and produce non-specific transcriptomic effects that confound interpretation of signaling responses, making it imperative that the system operate at modest expression levels. Existing dCas12a-based CRISPRi systems, however, operate reliably only under high expression and with short poly-crRNA arrays, owing to three compounding limitations: suboptimal effector performance^15^, competition among crRNAs for a limited pool of dCas12a^10^, and inefficient array processing^16^. We therefore reasoned that jointly optimizing intrinsic effector potency, component stability, and crRNA array architecture would be necessary to achieve scalable programmable repression of multiple genes (**Fig. 1C**, bottom).

To identify a dCas12a-effector variant with enhanced intrinsic potency, we generated a library of approximately 50 rationally designed variants. Each library member incorporated one of four dCas12a variants (LbCas12a, AsCas12a, hyperdCas12a^17^, and enAsCas12a^18^), three plasmid configurations, three epigenetic effector domains, four nuclear localization sequences, and used one of two possible domain arrangements (**Fig. 1D**). To systematically characterize repression potency, persistence, and specificity of the variants, we engineered a reporter cell line carrying a genomically-integrated destabilized mCitrine fluorescent protein constitutively expressed from an EF1α promoter. We utilized a compact, single-vector design that co-encodes the dCas12a-effector and the crRNA array (hereafter, the all-in-one construct). Transient delivery of individual library members into the reporter cell line induced epigenetic silencing of the EF1α promoter, rapidly reducing fluorescence from the destabilized mCitrine reporter, as measured by flow cytometry (**Fig. 1E**).

The screen established design rules for an effective dCas12a-effector variant. First, the Cas12a variant hyperdCas12a, an LbCas12a mutant with enhanced DNA binding^17^, consistently outperformed existing AsdCas12a variants in mediating transcriptional repression (**Fig. S1A**). Second, crRNA abundance is a major limiting factor for knockdown efficiency (**Fig. S1B**). Third, fusing the ZIM3-KRAB effector to the N-terminus of dCas12a results in stronger repression than C-terminal fusions (**Fig. S1C**). These insights guided the development of an optimized Cas12a variant, comprising a CAG-driven hyperdCas12a with an N-terminal fusion of ZIM3-KRAB, which we termed *ZK-dCas12a* (**Fig. 1F**). This variant surpassed existing dCas12a-effector variants in the speed, magnitude, and durability of repression of the mCitrine reporter (**Fig. 1F**) with no reduction in specificity (**Fig. S1D-E**). Whole-transcriptome RNA sequencing of cells transfected with this variant targeting the EF1α promoter showed a 5-fold decrease in mCitrine expression across biological replicates compared to a negative control transfected with non-targeting crRNA (**Fig. 1G**). The downregulation of mCitrine exceeded the differential expression of nearly all non-targeted genes (13628 out of 13631 genes, with the exceptions of DLEU7 and LOC105371271). Taken together, these results identify a dCas12a-effector variant and expression construct design with enhanced potency, persistence, and specificity over existing systems.

### ZK-dCas12a enables potent, specific, and sustained multiplexed repression of cell surface proteins

We next asked whether the new dCas12a-effector variant could achieve simultaneous knockdown of multiple endogenous targets. As a tractable model system, we selected CD surface proteins, whose turnover rates are comparable to those of TGF-β receptors^19,20^ and whose expression levels can be quantitatively assessed using flow cytometry with readily available antibodies.

To assess knockdown efficacy across a range of target abundances, we selected a panel of CD surface proteins spanning a broad range of basal expression levels in HEK293T cells (**Fig. S2A**). Transient delivery of the all-in-one constructs encoding either individual crRNAs or poly-crRNA arrays targeting these CD markers (**Fig. 2A**) resulted in >70% knockdown of all four targets (**Fig. 2B)**. Notably, the repressive activity of a given crRNA was not compromised within a poly-crRNA array (**Fig. 2B**), a phenomenon observed with previous dCas12a-based CRISPRi systems^10^.

**Fig. 2.**
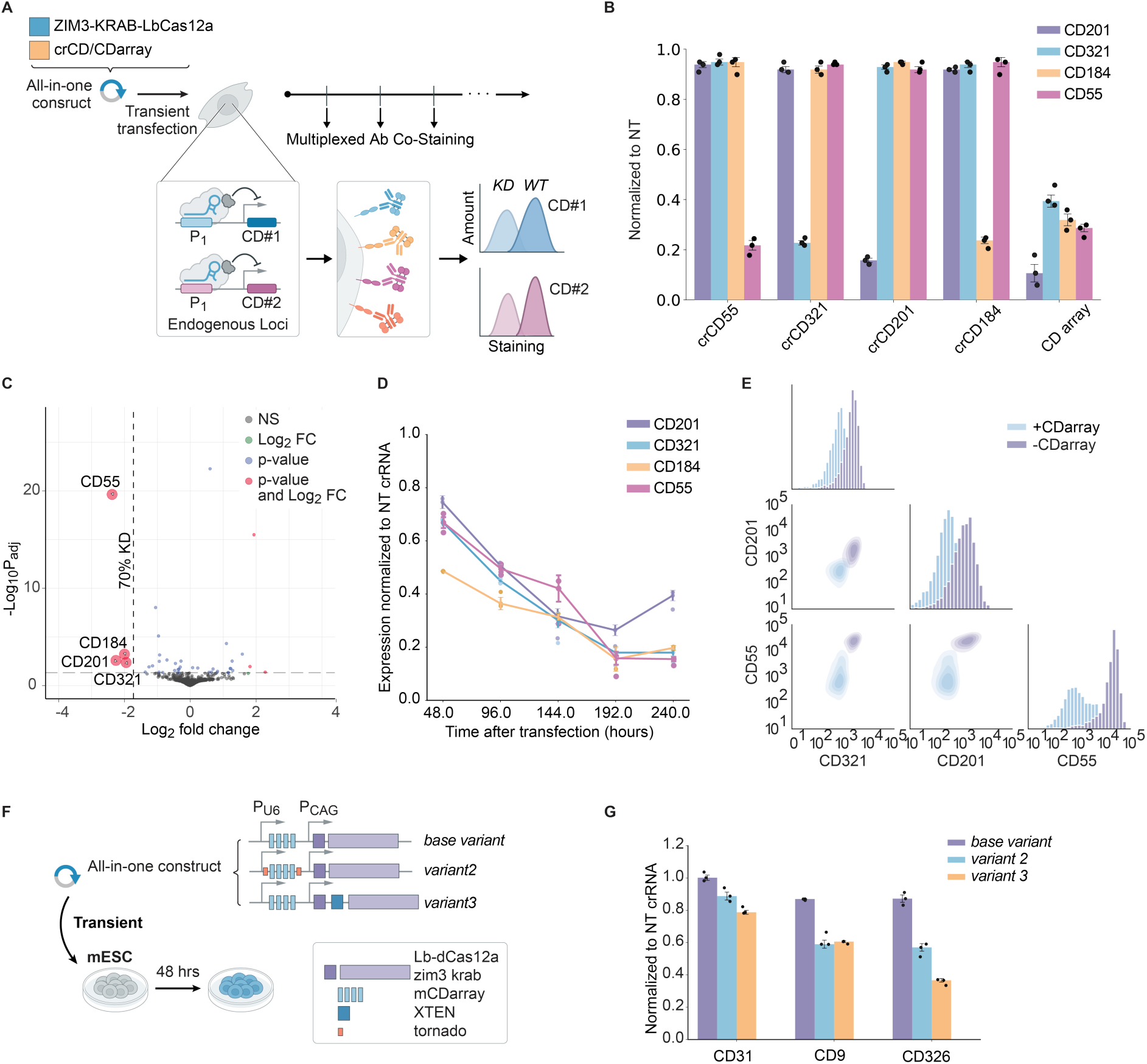
ZK-dCas12a achieves potent, specific, and sustained same-cell multiplexed perturbation of cell surface proteins. **(A)** Experimental workflow for multiplexed assessment of protein-level perturbation using ZK-dCas12a. Wild-type HEK293T cells were transiently transfected with the all-in-one construct co-expressing a poly-crRNA array targeting four CD surface proteins as well as the enhanced dCas12a-effector variant. Cells were subject to surface antibody (Ab) co-staining at varying time points after the initial transfection. **(B)** Flow cytometric measurement of CD marker surface levels 144 hrs after transfection. x-axis label indicates perturbation conditions. Dot denotes median staining intensity normalized to conditions transfected with non-targeting crRNA (NT crRNA) from an independent biological replicate. Bar heights and error bars denote the mean ± s.e.m. across three independent biological replicates. **(C)** Differential expression analysis of three independent RNA-seq experiments comparing samples perturbed with CD array vs NT crRNA. Dots denote individual genes. Genes with p_adj_ < 5x10^-4^ and fold-change > 1.73 are highlighted with red circles. Targets of CD array, CD55, CD184, CD201, and CD321 are highlighted with enlarged red circles. **(D)** Flow cytometric measurement of CD protein level at varying time points after initial transfection. Dot denotes median staining intensity normalized to conditions transfected with NT crRNA from an independent biological replicate. Bar heights and error bars denote the mean and s.e.m. across three independent biological replicates. **(E)** Pairplot analyses of flow cytometry result from CD surface marker staining. Diagonal histograms depict staining distributions of three examined CD markers. Off-diagonal contour plots are generated by kernel density estimations of individual single-cell pairwise measurements between all possible pairs of examined CD surface markers. **(F)** Schematic of the designs of different ZK-dCas12a constructs with stability-enhancement features. **(G)** Multiplexed flow cytometric measurements of mouse CD protein knockdown in mESC cell lines transiently transfected with different stability-enhanced versions of ZK-dCas12a. Dot denotes median staining intensity normalized to cells transfected with NT crRNA from independent biological replicates. Bar heights and error bars denote the mean ± s.e.m. across three independent biological replicates.

To assess specificity, we performed RNA sequencing on cells transfected with the poly-crRNA construct. The four targeted CD markers were the only genes significantly downregulated across the transcriptome (**Fig. 2C**). To evaluate the persistence of multiplexed gene repression, we performed longitudinal flow cytometry to track the expression of targeted CD proteins following the initial transient delivery of the poly-crRNA construct (**Fig. 2A**). Repression was sustained for at least ten days post-transfection (**Fig. 2D**). Moreover, examination of pair-wise relationships among targeted CD markers showed that the poly-crRNA construct achieved simultaneous knockdown of all CD markers at the single-cell level (**Fig. 2E**). Taken together, these results showed that transient delivery of the enhanced dCas12a-effector variant with a poly-crRNA array could achieve potent, specific, and sustained same-cell multiplexed repression of surface protein targets across a broad expression range.

Since ZK-dCas12a functions via endogenous epigenetic machinery, its performance could potentially vary across cell types. We therefore performed multiplexed knockdowns of the same set of CD proteins across diverse human cell types, as well as of an additional panel of mouse CD proteins (**Fig. S2B**) in mouse cells. Knockdown efficiencies were high (>50% protein reduction) across these cell types (**Fig. S3A**). Additionally, ZK-dCas12a achieved strong knockdown (ranging from 40% - 80% protein reduction) when transiently delivered into primary bone marrow cells (**Fig. S3B**). These results showed that ZK-dCas12a functions across multiple cellular contexts.

### Single-copy genome integration of stability-enhanced ZK-dCas12a enables uniform, synchronous, and tunable multiplexed genetic perturbation

Establishing defined receptor profiles and quantitatively measuring their signaling responses requires precise control of expression levels, which is difficult to achieve through the heterogeneous process of transient transfection. Site-specific genomic integration is ideal for signaling analyses as it ensures uniform perturbation, low cytotoxicity, and a defined genetic background.

However, the single-copy nature of site-specific genomic integration imposes a stringent expression ceiling, which historically limits the performance of dCas12a-based CRISPRi systems^17^. To test the performance of ZK-dCas12a in a low-expression regime characteristic of single-copy genomic integration, we transfected it into mouse embryonic stem cells, whose rapid doubling causes strong dilution of intracellular components, approximating the low steady-state concentrations of single-copy expression. In this regime, the *base variant* co-expressing ZK-dCas12a and a poly-crRNA array targeting mouse CDs substantially underperformed (**Fig. 2, F and G**), indicating that potency alone is insufficient when component levels are inherently constrained.

We reasoned that stabilizing the guide RNAs and effector protein could increase their activity. crRNAs are prone to degradation by endogenous exonucleases, which limits their steady-state intracellular concentration^21–23^. Circularizing RNA could mitigate this degradation by protecting against exonuclease activity^24,25^. We flanked the poly-crRNA array with the Tornado system—a ribozyme-based, self-circularizing RNA platform—to promote crRNA stabilization^26^. In parallel, we increased dCas12a protein stability by incorporating the XTEN linker, a peptide known to prolong protein half-life^27^. This approach has been successfully used in Cas9-based CRISPRi systems and recently adapted for Cas12a^10,28^. Finally, we incorporated these modifications into the all-in-one construct and generated two stability-enhanced variants: *variant 2*, with the Tornado system stabilizing crRNAs, and *variant 3*, with the XTEN-linker stabilized dCas12a (**Fig. 2F**). As anticipated, stability-enhanced versions (*variant* 2 and *variant* 3) increased repression by 20%–40% (**Fig. 2G**). Repression of CD31 was not as strong as that of other CD proteins, likely due to its high basal expression level. Nevertheless, taken together, these results showed that stability enhancement at both the protein and crRNA level allow ZK-dCas12a to perform effectively even at restricted expression level, preparing it for the constrained-expression regime of site-specific genomic integration.

We integrated a doxycycline (Dox)-inducible, stability-enhanced ZK-dCas12a cassette at the AAVS1 safe-harbor locus by CRISPR-HDR^29^ (**Fig. 3A**). This design combined four features: safe-harbor integration for a uniform, synchronous isogenic population; promoter-less targeting that couples integration with resistance to enable rapid clone derivation^30^; a bi-directional promoter to minimize transcriptional crosstalk in Dox-inducible systems^31^; and stability enhancement to preserve circuit performance at single-copy expression.

**Fig. 3.**
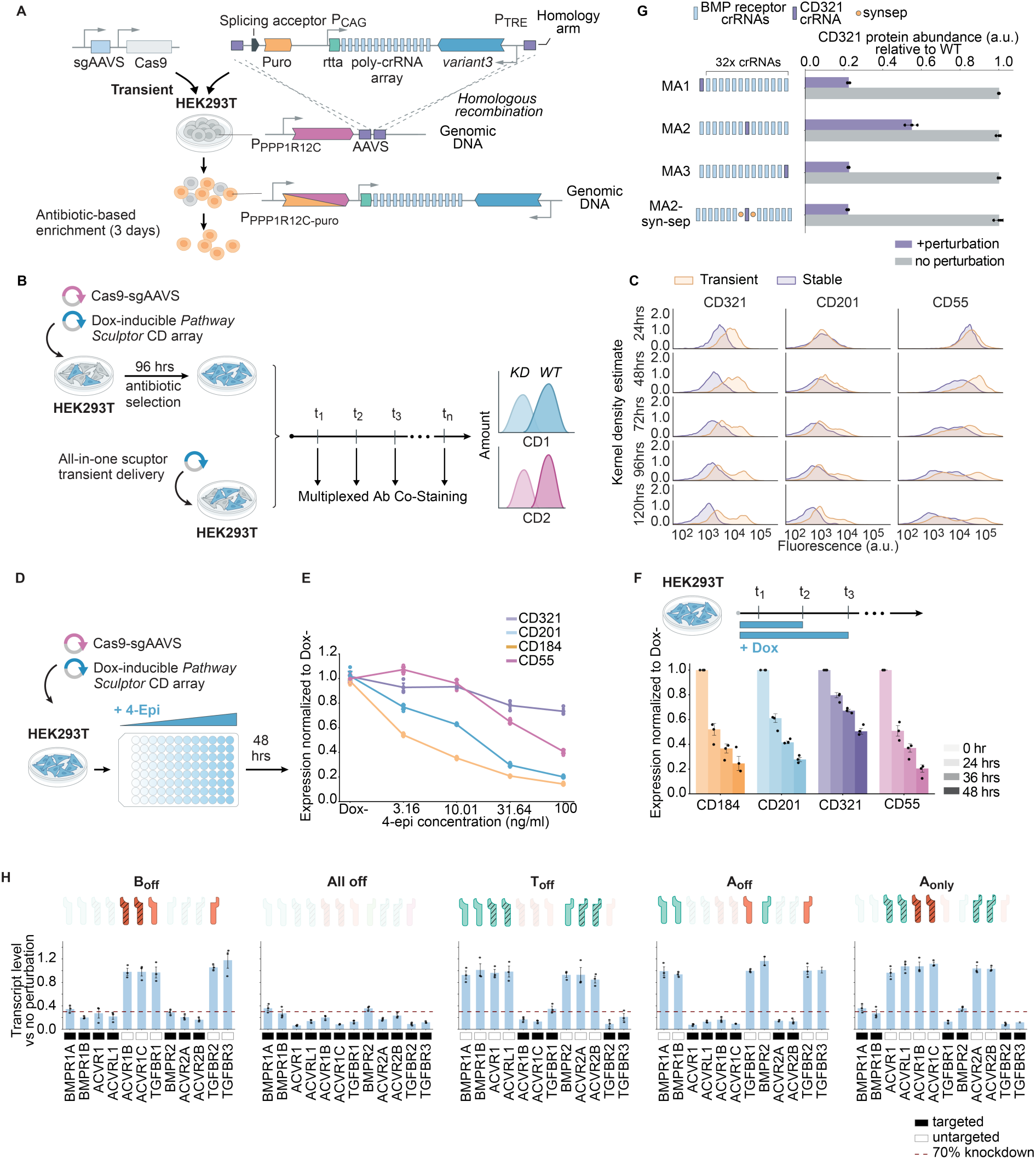
Engineered *Pathway Sculptor* cell line enables uniform, synchronous, and tunable multiplexed genetic perturbation. **(A)** Schematic of stable ZK-dCas12a cell line engineering: safe-harbor locus site directed targeting of Dox-inducible ZK-dCas12a construct. A donor plasmid with a splicing acceptor, selection cassette, poly-crRNA array, and ZK-dCas12a is co-transfected with a plasmid encoding Cas9 and AAVS1-sgRNA, and integrated into the human AAVS1 safe-harbor locus via homologous recombination, followed by antibiotic-based enrichment for 3 days. (**B**) Experimental workflow for comparing perturbation uniformity between transient delivery (all-in-one construct) and the engineered stable ZK-dCas12a cell line. Both systems target a CD surface protein array in HEK293T cells; uniformity is assessed by multiplexed antibody co-staining and flow cytometry at multiple time points. **(C)** Flow cytometric measurement of CD protein level at varying time points after Dox induction or after initial transfection of all-in-one construct, in the case of engineered stable cell line and wild-type cells, respectively. The y-values represent kernel density estimates of the probability density function. **(D-F)** Two titration strategies deployed in engineered ZK-dCas12a cell lines. Top: titratability is achieved through administration of different concentrations of Dox, or 4-ED. Bottom: titratability is achieved through administration of a saturated concentration of Dox for different durations of time. (**G**) Assessment of crRNA array processivity using MA1, MA2, MA3, and MA2+syn-sep variants. CD321 knockdown efficiencies are shown for each array variant, with (lavender) and without perturbation (gray). Incorporation of synthetic separators (syn-sep) rescues processivity in the MA2 configuration. **(H)** *Pathway Sculptor* can be used to generate diverse TGF-β superfamily receptor profiles. Transcript-level knockdown in cells carrying stable Dox-inducible *Pathway Sculptor* with poly-crRNA array targeting arbitrary subset or full set of twelve receptors, denoted by names and corresponding subunit cartoons. Black and white squares indicate targeted and untargeted receptor subunits. Maroon dashed line indicates 70% knockdown. Each dot represents an independent biological replicate; bar heights and error bars denote mean ± s.e.m.

The engineered cell line achieved more homogeneous expression than transient transfection. We compared the kinetics and uniformity of CD protein repression over 120 hours (**Fig. 3B**, top) following Dox induction against a transient delivery experiment using an all-in-one construct (**Fig. 3B**, bottom). In both settings, we observed progressive loss of all three targeted CD proteins (**Fig. 3C**). However, at each time point, the stable cell line exhibited more homogeneous marker repression than the transiently transfected cells (**Fig. 3C**). Thus, the engineered ZK-dCas12a cell lines provide uniform and synchronous multi-gene repression.

The ZK-dCas12a cell lines provided three tuning mechanisms: (1) modulating protein and/or crRNA levels (concentration tuning), (2) varying the duration of repression (duration tuning), and (3) incorporating mismatches into crRNAs to modulate binding strength (affinity tuning). To achieve concentration tuning, we treated polyclonal HEK293T cells with increasing concentrations of Dox or its analog, 4-epidoxycycline. While Dox induced all-or-none expression of ZK-dCas12a, 4-epidoxycycline allowed robust titration of ZK-dCas12a across two orders of magnitude (**Fig. S4A**). To different extents, this titration achieved graded knockdown of the four targeted CD markers (**Fig. 3D-E**). CD55, CD184, and CD201 exhibited a tunability range of up to 5-fold, whereas CD321 showed a more modest 1.5-fold reduction, likely due in part to its high basal expression. Tuning the duration of Dox exposure similarly led to graded repression across all markers (**Fig. 3F**), with CD321 again showing a more limited response. For affinity tuning, we introduced single mismatches at different positions of an EF1α-targeting crRNA to modulate its binding strength. This approach enabled fine-tuned repression of a mCitrine reporter upon transfection into the engineered mCitrine reporter HEK293T cell line (**Fig. S4B**). Together, these results demonstrate that engineered ZK-dCas12a cell lines support tunable multi-gene regulation with flexible and orthogonal control mechanisms.

### *Pathway Sculptor* reprograms TGF-β superfamily receptor profiles

We next asked whether the optimized ZK-dCas12a system could knock down an entire set of receptors and thereby enable controlled programming of receptor expression profiles. We first focused on the seven canonical BMP receptors. We adopted a multi-crRNA strategy, targeting the promoter of each gene by multiple crRNAs. This circumvents the trial-and-error typically required to identify accessible promoter binding sites and potentially allows synergistic enhancement of repression^32^. The multi-crRNA design resulted in a poly-crRNA array harboring a total of 32 crRNAs (1.3 kb), with each receptor targeted by 2–5 crRNAs.

The lengths of this and other arrays in this work exceed those of typical CRISPRi arrays. Previously, long arrays have encountered limitations in Cas12a-based systems^16,33,34^. Therefore, we asked whether similar issues might functionally inhibit the system. We designed three array variants—MA1, MA2, and MA3—each consisting of the full 32-crRNA BMP receptor-targeting array, with an additional crRNA inserted at different positions (**Fig. 3G**). The MA1 and MA3 stable cell lines, which incorporated CD321 crRNA at early and late positions, showed comparable CD321 knockdown, suggesting array transcription is preserved (**Fig. 3G**). MA2, with the CD321 crRNA in the middle, achieved knockdown, but to a lesser extent, indicating that processing efficiency varies with position. To overcome this, we incorporated 4-bp synthetic separators (syn-sep) to facilitate array processing by providing a favorable sequence context for Cas12a’s pre-crRNA cleavage activity^35^. An array incorporating syn-sep on both sides of CD321 crRNA (MA2-syn-sep) restored its CD321 knockdown efficiency to levels comparable to MA1 and MA3 (**Fig. 3G**), suggesting that incorporation of syn-sep sequences can overcome position-dependent processing limitations.

Turning to TGF-β, we next engineered a cell line with single-copy integration of stability-enhanced ZK-dCas12a and an optimized array targeting all seven BMP receptors. In this cell line, Dox induction achieved potent (>70%) transcript-level reduction of all BMP receptors (**Fig. 3H**, first panel), demonstrating that syn-sep incorporation enhances multiplexed repression in poly-crRNA arrays.

We further extended *Pathway Sculptor* to the entire TGF-β superfamily receptor repertoire. While the twelve canonical TGF-β superfamily receptors (seven type I and five type II) are all expressed in HEK293 cells (**Fig. S4C**), we made two curation decisions to focus the target set on receptors relevant to BMP/TGF-β signaling. First, we excluded AMHR2, whose ligand specificity is largely restricted to anti-Müllerian hormone^36^ and whose engagement is therefore orthogonal to the BMP and TGF-β branches examined here. Second, we added TGFBR3 (betaglycan), a type III co-receptor that lacks intracellular kinase activity but functions as a critical accessory factor for TGF-β ligand presentation and access to the type I/II receptor complex^37^. The resulting target set comprises eleven canonical type I/II receptors (the original twelve minus AMHR2) plus TGFBR3, again totaling twelve receptors. To target this set simultaneously, we expanded the poly-crRNA array to 52 crRNAs, and integrated this construct at the AAVS1 safe-harbor locus as above. Remarkably, Dox induction achieved strong, simultaneous repression of all twelve receptors (**Fig. 3H**, second panel).

To produce defined receptor profiles, we developed a time- and cost-effective subset-removal strategy that excises selected crRNAs from the full array while preserving the rest (**Fig. S5B**). Applying this strategy, we generated four configurations spanning the major axes of the receptor network: B_off_ (the seven-BMP-branch-receptor knockdown introduced above), T_off_ (the five TGF-β-branch receptors knocked down, including TGFBR3), A_off_ (the six ACVR-class receptors knocked down, with the five non-ACVR BMPR/TGFBR and TGFBR3 retained), and A_only_ (the six non-ACVR receptors knocked down, with all six ACVRs retained). Each configuration achieved potent and specific receptor depletion (**Fig. 3H**, third to fifth panel), yielding a panel of programmable receptor expression profiles.

Together, these results showed that joint engineering of array architecture, together with the more potent dCas12a-effector (**Fig. 1**) and its stability enhancement (**Fig. 2**), culminated in a significantly improved dCas12a-CRISPRi system enabling programmable reconfiguration of pathway profile. Thus, we termed the system *Pathway Sculptor*.

### BMP and TGF-β branches of the TGF-β superfamily pathway exhibit cell-intrinsic, bidirectional crosstalk

So-called crosstalk between the BMP and TGF-β branches plays important roles across diverse pathophysiological contexts^2,38,39^. However, it remains unclear whether such crosstalk happens at the single-cell level, how strong it is quantitatively, and, more generally, whether it represents an intrinsic feature of the pathway architecture.

We first sought to establish a quantitative single-cell readout of both branches in wild-type cells. We engineered a HEK293 cell line carrying two genomically-integrated fluorescent reporters driven by SMAD1/5/8-responsive (BRE) elements and SMAD2/3-responsive (TRE) elements (**Fig. 4A**)^40,41^. Each reporter responded to its cognate ligands (intra-branch signaling) in a dose-dependent manner, with high signal-to-noise ratios at saturating doses (**Fig. 4B**, first and fourth row). Consistent with previous work, some ligands, including TGF-β1 and TGF-β2, showed biphasic responses^42–44^. These biphasic responses also occurred in the BRE responses to BMP10, BMP4, and BMP5 (**Fig. 4B**, first row). Together, these results showed that the dual-reporter cell line enables quantitative analysis of signaling transduction through the canonical BMP and TGF-β signaling branches at the single-cell level.

**Fig. 4.**
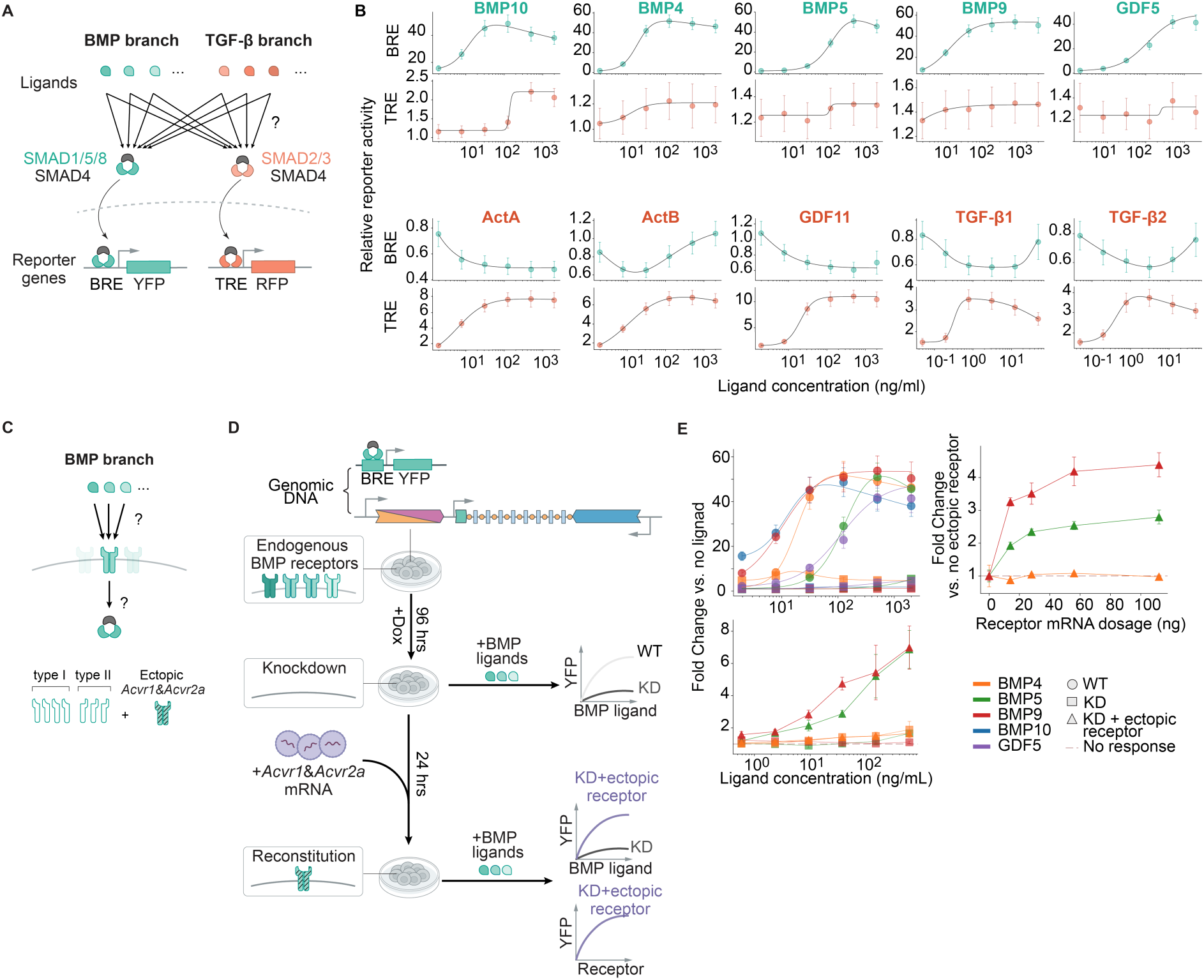
The TGF-β superfamily exhibits bidirectional crosstalk while individual receptor pairs mediate ligand-selective responses. (**A**) Schematic of the dual-reporter system for simultaneous measurement of BMP and TGF-β branch signaling. BMP-family ligands (green) and TGF-β-family ligands (red) activate the SMAD1/5/8–SMAD4 and SMAD2/3–SMAD4 transducers, respectively. Genomically-integrated BRE-YFP reporter and TRE-RFP reporter are used to read out BMP branch and TGF-β branch signaling activities, respectively. (**B**) Dose-response curves for ten TGF-β superfamily ligands. Odd rows (green): BRE reporter activity; Even rows (red): TRE reporter activity. Points and error bars denote mean ± s.d. across biological replicates; solid lines denote fitted dose-response curves using either Hill or biphasic models. (**C**) Schematic illustrating the BMP receptor reconstitution strategy. Ectopic delivery of mouse *Acvr1* and *Acvr2a* mRNA into the receptor-depleted cells reconstitutes Acvr1-Acvr2a receptor pairs to allow quantitative assessment of the dependencies of its signaling capabilities on ligand dosages and receptor expression levels. (**D**) Experimental workflow for BMP receptor reconstitution and its functional assessment. (**E**) Top: BMP signaling responses to BMP10, BMP4, BMP5, BMP9, and GDF5 in no-perturbation (circles) versus *Pathway Sculptor*-induced knockdown (squares) conditions, measured as BRE-YFP fold change normalized to unstimulated controls across a range of ligand concentrations. Lines denote fitted dose-response curves. Middle: BMP signaling response restoration as a function of ligand dosage in receptor reconstituted condition (triangle) versus *Pathway Sculptor*-induced knockdown condition (squares). Bottom: BMP signaling response restoration as a function of mouse Acvr1/Acvr2a mRNA dosage (0–100 ng) for BMP5, BMP9, and BMP4. Fold changes are normalized to zero receptor dosage control. All error bars denote mean ± s.e.m from three independent biological replicates.

We next used the dual-reporter cell line to analyze inter-branch signaling, i.e. BRE responses to TGF-β ligands and TRE responses to BMP ligands. Surprisingly, all five BMP ligands tested exhibited weak, but dose-dependent, activation of the TRE reporter (**Fig. 4B**, second row). Conversely, the TGF-β ligands regulated the BRE reporter, but in more complex ways (**Fig. 4B**, third row). For example, ActA and GDF11 suppressed BRE activity in a dose-dependent manner, while ActB, TGF-β1, and TGF-β2 generated inverted biphasic responses compared to the corresponding TRE reporter. These results revealed cell-intrinsic, bidirectional inter-branch crosstalk, potentially mediated by competition for shared pathway components.

### A single receptor pair mediates differential signaling of distinct BMP ligands

We next asked whether different ligands can be distinguished by a single receptor pair, or whether discrimination requires the full repertoire of receptor variants. To address this, we devised a receptor reconstitution strategy to measure the signaling capabilities of individual ligand-receptor complexes, i.e. how SMAD activation responds to varying ligand concentrations and receptor expression levels (**Fig. 4C**). Using the B_off_ cell line (**Fig. 3H**) carrying a SMAD1/5/8 reporter, we depleted all BMP receptor subunits and replaced them with a single type I-type II receptor pair in HEK293 cells (**Fig. 4D**). Dox induction of *Pathway Sculptor* eliminated signaling responses to all five tested BMP ligand variants across a broad range of ligand concentrations (**Fig. 4E**, top left). Ectopically expressing mouse *Acvr1-Acvr2a* receptor pair in the knockdown condition restored signaling responses to BMP5 and BMP9 (**Fig. 4E**, bottom left), albeit to an order of magnitude lower than in wild-type cells. By contrast, signaling response to BMP4 remained negligible (**Fig. 4E**, bottom left). Notably, the signaling response of different ligands exhibited distinct dependencies on ectopic receptor expression (**Fig. 4E**, top right): while BMP5 and BMP9 responses increase monotonically with receptor dosage, BMP4 exhibits no dependence on receptor level. Together, these results demonstrated the ability to replace endogenous receptor families with defined receptor variants, and showed that a single receptor pair can mediate graded, ligand-selective responses in a manner tunable by receptor expression level.

### Intra-branch signaling requires off-branch receptors

A longstanding puzzle is how the BMP and TGF-β branches of the overall TGF-β superfamily interact. Do BMP ligands signal through TGF-β receptors, or *vice versa*? And, which sets of R-SMADs mediate such signaling (**Fig. 5A**)? To address these questions, we incorporated BRE/TRE reporters into previously established cell lines with alternative receptor profiles and analyzed their signaling responses (**Fig. 3H, Fig. S5A**).

**Fig. 5.**
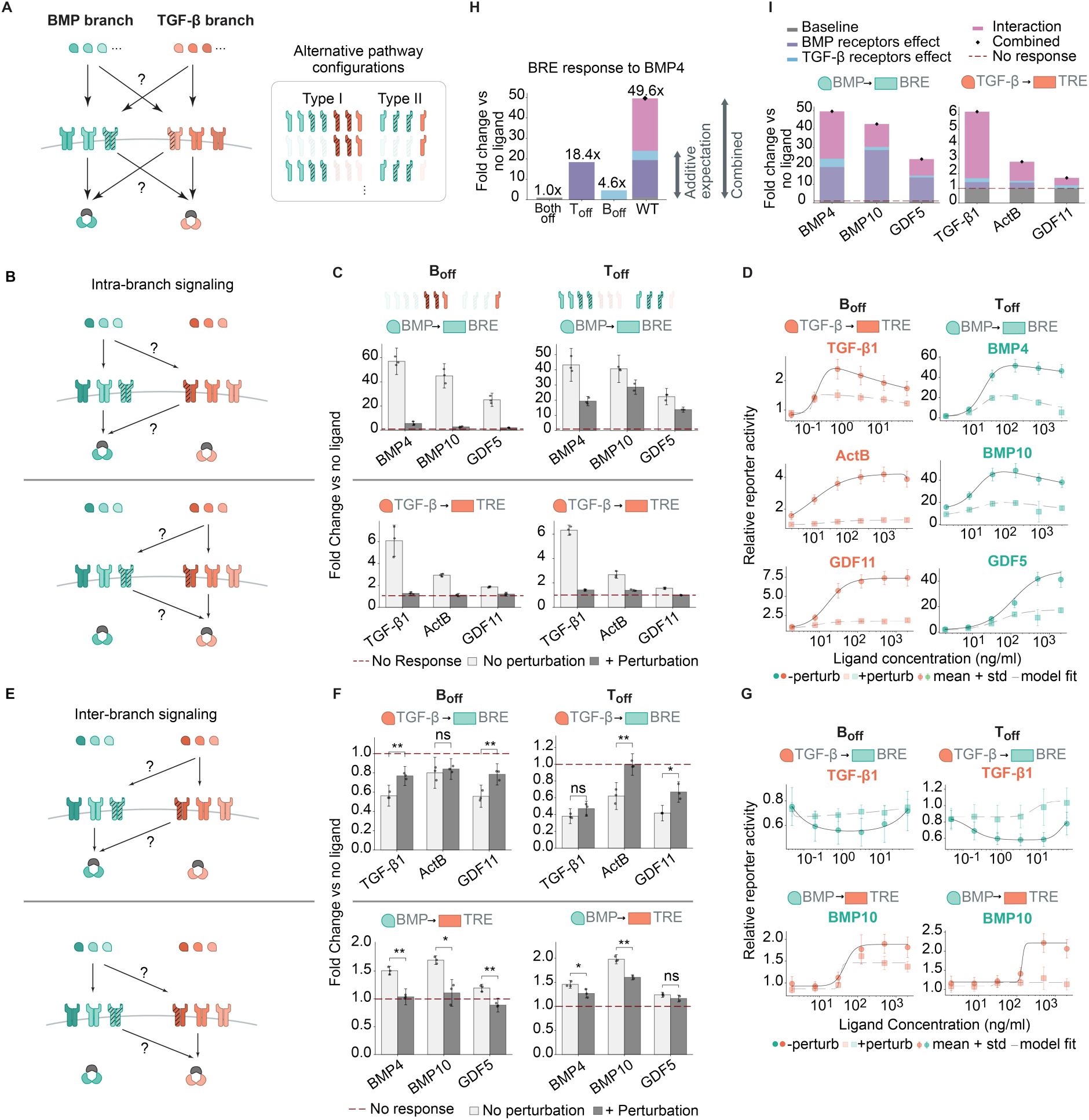
Programmable receptor profiles reveal functional interdependencies of the BMP and TGF-β branches. (**A**) Schematic of BMP and TGF-β signaling flows and engineered receptor configurations examined in this figure. (**B**) Putative signaling transduction paths through intra-branch signaling. (**C**) Intra-branch BRE-YFP (top) and TRE-RFP (bottom) responses to BMP-family ligands (BMP4, BMP10, GDF5) and TGF-β-family ligands (TGF-β1, ActB, GDF11) in B_off_ (left) and T_off_ (right) configurations. Fold change at saturating ligand concentration normalized to unstimulated control; dashed line: no-response baseline. (**D**) Representative dose-response curves for selected ligands in B_off_ (left) and T_off_ (right) configurations. Circles: +perturbation; squares: −perturbation. Points: mean ± s.d.; lines: fitted Hill or biphasic models. (**E**) Putative signaling transduction paths through inter-branch signaling. (**F**) Inter-branch BRE-YFP (top) and TRE-RFP (bottom) responses to TGF-β-family ligands (TGF-β1, ActB, GDF11) and BMP-family ligands (BMP4, BMP10, GDF5) in B_off_ (left) and T_off_ (right) configurations. Significance vs. no-response baseline by Welch’s t-test (** p < 0.01; ns, not significant). (**G**) Representative dose-response curves for inter-branch ligand responses. (**H**) Interaction decomposition of BRE response to BMP4. Bars show fold changes for Both off (no receptor), T_off_, B_off_, and wild-type conditions; stacked components indicate baseline, BMP receptor, TGF-β receptor, and interaction contributions. (**I**) Decomposition of BRE and TRE responses into additive receptor contributions and interaction terms across BMP- and TGF-β-family ligands. Stacked bars: baseline, BMP receptor, TGF-β receptor, cross-branch interaction, and combined effects.

We first analyzed intra-branch signaling in the B_off_ and T_off_ conditions (**Fig. 5, B and C**). These cell lines exhibited strongly reduced signaling in response to ligands associated with the knocked down receptor branch. As expected, B_off_ became non-responsive to the BMP ligands BMP4, BMP10, and GDF5 (**Fig. 5C**, upper left). Conversely, T_off_ lost responsiveness to TGF-β1, ActB, and GDF11 (**Fig. 5C**, bottom right).

More surprisingly, knockdown of off-branch receptors also inhibited intra-branch signaling. For example, the B_off_ condition exhibited a nearly complete loss of TRE responses to TGF-β ligands (**Fig. 5C**, bottom left). Conversely, the T_off_ condition exhibited diminished BRE responses to all three BMP ligands examined (**Fig. 5C**, top right). Notably, these effects were observed across a broad range of ligand dosages (**Fig. 5D**), suggesting that they could be physiologically relevant. Thus, receptors associated with the BMP and TGF-β branches are required not only for signaling by their own cognate ligands, but also for signaling by non-cognate ligands.

To determine whether these reciprocal receptor dependencies generalize beyond HEK293 to other cell types and species, we stably integrated the same dual TRE/BRE reporters into the NAMRU mouse mammary gland (NMuMG) epithelial cell line, which has been previously used to study BMP responses^45^. We then designed crRNA arrays targeting mouse BMP and TGF-β receptors. Finally, as above, we knocked down receptor subsets and analyzed responses to corresponding mouse ligands. These experiments revealed similar, although in some cases weaker, reciprocal dependencies on off-branch receptors for intra-branch signaling (**Fig. S6A**).

Taken together, these results showed that BMP and TGF-β intra-branch signaling unexpectedly depends on receptors in the other branch, in both mouse and human cells.

### Inter-branch signaling requires receptors from both signaling branches

The finding that intra-branch signaling requires receptors from both branches raises an analogous question for inter-branch signaling: do BMP ligands require TGF-β receptors to activate the TRE reporter, and *vice versa*? To address this, we measured inter-branch signaling responses in the B_off_ and T_off_ conditions (**Fig. 5E**). Strikingly, both B_off_ and T_off_ exhibited modest, but significant, increases in BRE responses to some TGF-β ligands (**Fig. 5F**, top row), suggesting that removing receptors from either branch partially relieves competitive inhibition between the two branches. By contrast, TRE responses to BMP ligands were significantly reduced in both conditions (**Fig. 5F**, bottom row). Notably, these effects were broadly observed across a range of ligand concentrations (**Fig. 5G**). Together, these results demonstrated that inter-branch signaling, like intra-branch signaling, requires the concerted action of receptors from both branches.

### BMP and TGF-β receptors cooperatively mediate signaling

These reciprocal receptor dependencies provoke the question of whether different sets of receptors synergize or antagonize each other’s effects. *Pathway Sculptor* allowed us to isolate the contribution of each receptor family in cells lacking the remaining receptor variants, and to measure deviations of their combined effects from additivity. For example, focusing on the BRE response to BMP4 (**Fig. 5H**), TGF-β receptors alone (the B_off_ configuration) generated an approximately 4.6-fold increase relative to full receptor knockdown, while BMP receptors alone (the T_off_ configuration) generated an approximately 18.4-fold increase. Yet wild-type cells containing both branches exhibited a ∼50-fold response (**Fig. 5H**), exceeding the sum of the two single-branch conditions. Thus, BMP and TGF-β receptor sets synergize to mediate BMP4 signaling to the BRE. Strong synergies between the two receptor sets were also observed for all other tested ligands (**Fig. 5I**), and, to a lesser extent, for inter-branch signaling as well (**Fig. S6B**). Together, these results show that the BMP and TGF-β branch receptors act synergistically to mediate signaling responses to BMP and TGF-β ligands.

### ACVRs mediate inter-branch crosstalk

The observed synergy between BMP and TGF-β receptors could potentially arise through inter-family receptor complexes. Indeed, a growing body of evidence suggests the formation of chimeric receptor complexes comprising variants from both signaling branches, with most cases involving at least one member of the ACVR class^46–49^. However, it has remained unclear how these complexes affect signaling.

*Pathway Sculptor* uniquely allows dissecting the role of inter-family complexes through perturbations along the ACVR class axis (ACVR vs non-ACVR; **Fig. 1B**). We focused on A_off_ and A_only_ (**Fig. 3H**), which by construction are enriched for intra-family and inter-family receptor complexes, respectively (**Fig. S6C**). Both A_off_ and A_only_ conditions strongly reduced intra-branch signaling responses (**Fig. S6D**), consistent with the involvement of ACVRs (the A_only_ configuration) and BMPR/TGFBRs (the A_off_ configuration) in signaling through both branches. By contrast, inter-branch signaling responses were only reduced in the A_off_ condition, but not the A_only_ condition (**Fig. 6A**). These results showed that ACVRs mediate crosstalk between the BMP and TGF-β branches, presumably through inter-family receptor complexes.

**Fig. 6.**
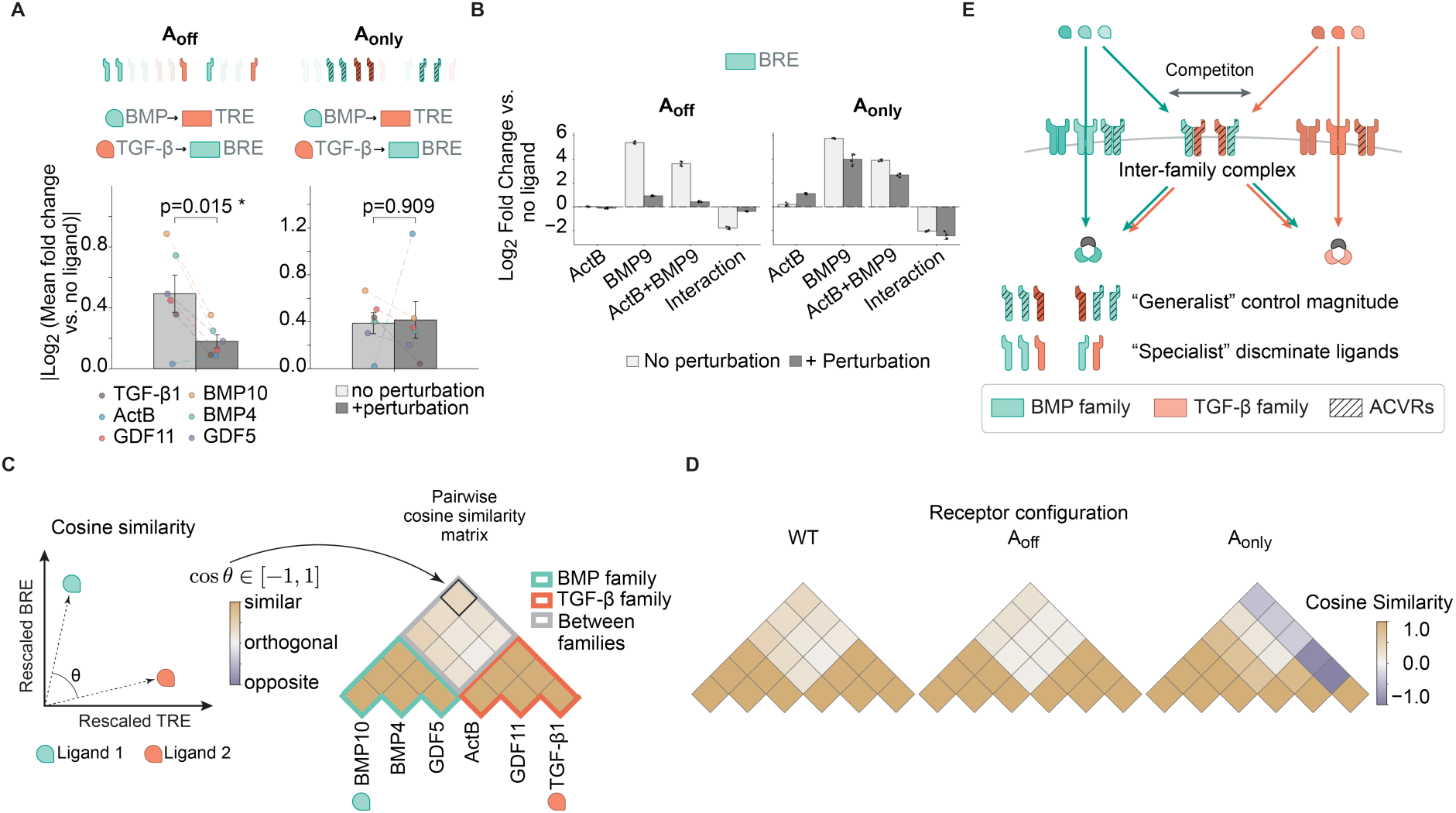
Shared ACVRs control signaling magnitude while branch-specific BMPRs/TGFBRs discriminate among ligands. (**A**) Aggregated BRE-YFP responses to TGF-β-family ligands and TRE-RFP responses to BMP-family ligands in A_off_ (left) and A_only_ (right) configurations. Significance vs. wild-type by Wilcoxon signed-rank test (A_off_: p = 0.015; A_only_: p = 0.909). Each dot represents the mean response of a given ligand across three biological replicates. (**B**) Log_2_-fold change of BRE-YFP responses in A_off_ (left) and A_only_ (right) to ActB alone, BMP9 alone, ActB + BMP9 combined, and the calculated interaction term. Light gray: wild-type; dark gray: A_off_ or A_only_. **(C)** Left: Schematic illustrating the cosine similarity metric. Each ligand’s signaling effect is represented as a vector of rescaled BRE-YFP and TRE-RFP responses; cosine similarity between two ligand vectors ranges from −1 (opposite) to 1 (similar), with 0 indicating orthogonality. Right: Pairwise cosine similarity matrix computed across all tested ligands in wild-type cells. Cosine similarity within the BMP family, within the TGF-β/activin family, and between families are highlighted with green, pink, and grey squares, respectively. **(D)** Pairwise cosine similarity matrices for WT (wild-type), A_off_, and A_only_ configurations. Color scale as in (C). **(E)** Schematic illustration of receptor functional specialization. BMPRs and TGFBRs only mediate intra-branch signaling and discriminate ligands; ACVRs mediate both intra- and inter-branch crosstalk without altering ligand discrimination; together, the receptor network shapes both signaling magnitude and ligand discrimination.

Inter-family complex formation could in principle impose receptor-level competition — a common mechanism underlying ligand antagonism. Ligand antagonism within the BMP family is prevalent^50^, and similar antagonism across families has also been observed. For example, ActB antagonizes BMP9 in diverse physiological contexts^51,52^. We therefore asked whether ACVRs could underlie ligand antagonism across families. To drive the system into a receptor-limited regime, we treated cells with ActB and BMP9 individually or in combination at saturating doses. ActB strongly antagonizes BRE reporter response to BMP9 (**Fig. 6B**), consistent with the model in which the two ligands compete for a shared pool of receptors. Strikingly, such antagonism was abolished in the A_off_ condition but remained unchanged, or slightly enhanced, in the A_only_ condition (**Fig. 6B**), suggesting that ACVRs serve as the primary source of competition at the receptor level. Taken together, these findings strongly suggest that ACVR-containing complexes govern inter-branch crosstalk within the TGF-β signaling network, likely through their capacity to form inter-family receptor complexes.

### BMPRs and TGFBRs play primary roles in ligand identity discrimination

A key function of any signaling system is to distinguish among different input signals. Given that ACVRs and non-ACVRs play distinct roles in routing signals within and across branches, we next asked whether they also play distinct roles in discriminating among ligands.

Because each TGF-β superfamily ligand elicits both TRE and BRE responses, ligand identity can be captured as a 2-D response vector. In wild-type cells, plotting these vectors showed that the two ligand families segregate cleanly along their respective axes (**Fig. S6E**), with consistent off-axis components reflecting the established crosstalk (**Fig. 4B**). To quantify discriminability, we used cosine similarity to compare every pair of ligand response vectors, yielding a similarity matrix across all ligands (**Fig. 6C**, schematic). In wild-type cells, cosine similarities are high within each family and low between families (**Fig. 6C**, matrix), confirming that this metric captures family-level ligand identity.

Interestingly, despite exhibiting substantially reduced intra- and inter-branch signaling magnitudes (**Fig. S6D** and **Fig. 6A**), the A_off_ condition minimally affected ligand discriminability, as evidenced by a cosine similarity profile nearly indistinguishable from those of wild-type cells (**Fig. 6D** and **Fig. S7**). By contrast, the A_only_ condition exhibited specific, ligand-selective breakdowns in discriminability (**Fig. 6D** and **Fig. S7**): ActB became more similar to BMP-family ligands, while TGF-β1 emerged as an isolated ligand not clearly associated with either family.

Taken together, these results reveal a functional division of labor within the TGF-β superfamily receptor network: ACVRs modulate signaling magnitude in a ligand-agnostic manner, while branch-specific BMPRs and TGFBRs discriminate among distinct ligand classes.

## Discussion

Metazoan cell-cell communication pathways are often built from families of paralogous, partially redundant, and broadly co-expressed components. This architecture allows cells to generate context-dependent responses, but it also makes it difficult to dissect information flow within them through conventional genetic approaches, where only a handful of genes can be simultaneously perturbed. What has been needed is the ability to programmably remove or modulate expression of arbitrary component combinations, including large subsets, within the same cell. In this study, we engineered *Pathway Sculptor*, a scalable multiplexed dCas12a-based CRISPRi platform, and applied it to dissect information flow within the TGF-β superfamily.

Building *Pathway Sculptor* revealed general engineering principles for achieving durable, potent, multiplexed epigenetic perturbation. Here, introduction of the ZIM3-KRAB effector domain, incorporation of stabilizing features to the protein, and enhancement of poly-crRNA processing together allowed *Pathway Sculptor* to achieve all three objectives simultaneously across a wide range of cellular contexts. Prior approaches, including siRNA, shRNA, miRNA, Cas9-based CRISPRi/a, and Cas13d, each optimized a subset of these performance features at the expense of others^28,53–59^. Additionally, with Cas13d, collateral RNA editing constrains its use across cell types^9^.

*Pathway Sculptor* provides a missing experimental handle for paralog-rich pathways such as Wnt, Notch, RTK, and cytokine signaling. In these pathways, it has long been possible to explore ligand combinations, but systematically accessing diverse receptor combinations has remained challenging. Now, arbitrary receptor profiles can be programmed and tested against different ligands or ligand mixtures, enabling a more complete combinatorial view of pathway function. For example, it should now be possible to determine how ligand-discrimination capacity scales with receptor diversity, or whether specific receptor profiles confer ratio-, fold-change-, or absolute-concentration sensing. More broadly, *Pathway Sculptor* is not restricted to single protein families: it allows simultaneous perturbation of arbitrary gene sets — for example, members of a transcription factor family, mixed combinations of regulators across pathways, or panels of chromatin modifiers. Its modularity also opens avenues for further engineering. For example, fusing transcriptional activation domains to *Pathway Sculptor* variants with orthogonal array binding preferences would enable bidirectional (activation as well as repression), multi-locus modulation in the same cell.

Using *Pathway Sculptor,* we systematically analyzed the signaling behaviors of a wide variety of TGF-β receptor profiles. These experiments revealed ligand-dependent crosstalk between the BMP and TGF-β branches (**Fig. 6E**). Unexpectedly, intra-branch signaling depended strongly on receptors from the other branch: removing BMP-branch receptors disrupted TGF-β-branch signaling, and *vice versa* (**Fig. 5**). Thus, receptors are functionally interconnected and cannot be neatly segregated into two independent families. This is supported here by the ACVR-specific loss of inter-branch responses (**Fig. 6A**) and ACVR-dependent ligand antagonism (**Fig. 6B**). Mechanistically, such crosstalk is consistent with prior reports of ACVR-containing interfamily receptor complexes^46–48^. The ACVR axis represents a distinct mode of receptor-mediated control orthogonal to the conventional BMP vs TGF-β dichotomy. Moreover, this work identifies a functional division of labor in the receptor network: the “generalist” nature of the ACVR receptors allows them to modulate signaling magnitude in a ligand-agnostic manner without substantially altering ligand discrimination (**Fig. 6D**). In contrast, the “specialist” receptors, BMPRs and TGFBRs, primarily convey signaling within their cognate branches, and therefore play a dominant role in discriminating among distinct ligands. Together, these findings suggest that TGF-β superfamily signaling operates as an integrated network, in which interfamily receptor complex formation and competition for shared receptors collectively shape signal propagation, pathway crosstalk, and ligand interpretation. Notably, recent work has shown that mixed SMAD1/5/8–SMAD2/3 complexes form under co-stimulation with BMP4 and TGF-β1 and are sufficient, in a mathematical model, to explain asymmetric crosstalk between TGF-β1 and BMP4 in NMuMG cells^60^. The results here are broadly consistent with this work, but suggest that effects on the composition of SMAD trimers could reflect receptor subunit composition.

A defining, but historically vexing, feature of TGF-β biology is its cell- and tissue-context dependence. For example, TGF-β1 suppresses early-stage epithelial tumors but drives invasion and metastasis in advanced disease^61^, while BMPR2 loss-of-function causes pulmonary arterial hypertension but is tolerated in other tissues^62^. Our results suggest that receptor expression profiles could produce context dependent effects. Because the BMP and TGF-β branches form a single integrated network, signaling by a given ligand or ligand combination can in general depend on the full receptor profile a cell expresses, which varies widely across cell types^1^. The asymmetric receptor architecture (**Fig. 6D**) gives cells distinct receptor-level handles for tuning this response: a cell could modulate signaling sensitivity by adjusting ACVR expression while preserving ligand discrimination. By contrast, modulating BMPR/TGFBR expression could reshape ligand-specificity. Receptor-level response tuning could help explain mechanistically why the same ligand can have divergent consequences in different tissues. Component multiplicity is a ubiquitous feature of most biological systems. By allowing same-cell multiplexed perturbations, *Pathway Sculptor* should enable researchers to analyze a broader repertoire of physiologically relevant pathway configurations than previously feasible. Such analyses may reveal principles of combinatorial control that were previously obscure.

## Acknowledgements

We thank Xiaojing Gao, Eliezer Calo, Dhiraj Indana and Judy Shon for comments on the manuscript; Yodai Takei, Lukas Moeller, Andrew Lu for advice on experimental design and technical support; Dongyang Li for scientific inputs; Inna-Marie Strazhnik for graphical design; Leah Santat, Jo Leonardo, and Rui Malinowski for administrative support. This research was supported by NIH R01EB030015 and NIH U01DK127420. M.B.E. is a Howard Hughes Medical Institute Investigator. B.G. is supported by the Damon Runyon Fellowship (DRG2441-21). G.M. is supported by the Human Frontiers Science Program (LT0037/2023-L). J.G. is supported by the Boehringer Ingelheim Funds PhD fellowship. B.A. and R.L. were supported by NIH R35HL150826 and R01DK143671. The content is solely the responsibility of the authors and does not necessarily represent the official views of the National Institutes of Health.

## Author contributions

B.G. and M.B.E. conceived and designed the study. M.B.E. directed and supervised the study. B.G., J.M.L., and M.B.E. directed and supervised experiments corresponding to Fig. 5 and Fig. 6. G.M., J.G. and R.H. assisted with or offered guidance regarding experiments corresponding to Fig. S3A. B.A. performed bone marrow knockdown experiments in Fig. S3B. H.L. assisted with experiments corresponding to supplementary figures. B.G.H. assisted with molecular cloning and longitudinal flow cytometry experiments. B.G. analyzed data. B.G. and M.B.E. wrote the manuscript with input from all co-authors.

## Declaration of generative AI and AI-assisted technologies in the manuscript preparation process

During the preparation of this work the author(s) used GitHub Copilot in Visual Studio Code in order to generate plotting codes. After using GitHub Copilot in Visual Studio Code, the author(s) reviewed and edited the content as needed and take(s) full responsibility for the content of the published article.

## Data and code availability

Raw and processed RNA-seq data have been deposited at the NCBI Gene Expression Omnibus under accession: GSE327757. Raw data and custom analysis code is available from the lead contact upon request.

## STAR Methods

### Tissue culture and cell lines

HEK293T cells were cultured in DMEM (Thermo Fisher Scientific) + 10% FBS (Avantor® Seradigm Research Grade Fetal Bovine Serum lot #89510-190) + Pen/Strep/L-glutamine and passaged every 2-3 days.

U2OS cells were cultured in DMEM (Thermo Fisher Scientific) + 10% FBS (Avantor® Seradigm Research Grade Fetal Bovine Serum lot #89510-190) and passaged every 2-3 days.

K562 Cells were grown in RPMI medium 1640 (Thermo Fisher Scientific) + GlutaMAX (Thermo Fisher Scientific) + 10% heat inactivated FBS (Invitrogen) + Pen/Strep (Thermo Fisher Scientific) + L-glutamine (Thermo Fisher Scientific) + sodium pyruvate (Thermo Fisher Scientific). Cells were maintained at the density of around 1 x 10^6^ cells/ml and passaged every 2-3 days.

E14 mES cells (ATCC cat. No. CRL-1821) were cultured in medium containing DMEM (Sigma), 15% ES cell qualified FBS (Gibco), 1x MEM non-essential amino acids (Thermo Fisher Scientific), 1 mM sodium pyruvate (Thermo Fisher Scientific), 100 μM β-mercaptoethanol (Thermo Fisher Scientific), 1x penicillin-streptomycin-L-glutamine (Thermo Fisher Scientific) and 1000 U/mL leukemia inhibitory factor (Millipore). Cells were maintained on polystyrene (Falcon) plates coated with 0.1% gelatin (Sigma) at 37 °C and 5% CO_2_ and passaged every 48 hrs.

NMuMG cells (NAMRU mouse mammary gland epithelial; ATCC CRL-1636) were cultured in DMEM (high Glucose) (Thermo 11965092) + 10% FBS (Avantor Seradigm) + NEAA (Thermo 11140050) + Sodium Pyruvate (Thermo 11360070)+ 1× Pen/Strep/L-glutamine and passaged every 3–4 days using 0.25% trypsin-EDTA. Cells were maintained below 80% confluency.

All cells are routinely tested for mycoplasma contamination.

### dCas12a-effector library design and screening

A library of approximately 50 dCas12a-effector variants was designed by combinatorially varying five parameters: (i) Cas12a backbone (LbCas12a, AsCas12a, hyperdCas12a, and enAsCas12a), (ii) promoter (CAG, EF1α, PGK), (iii) epigenetic effector domain (ZIM3-KRAB, KOX1-KRAB, Tandem-KOX1-KRAB), (iv) nuclear localization sequence (SV40-NLS, bipartite SV40, c-Myc-NLS), and (v) effector–dCas12a domain arrangement (N-terminal vs. C-terminal fusion). Each variant was assembled by Gibson cloning into an all-in-one backbone co-encoding a U6-driven crRNA cassette and verified by whole-plasmid Nanopore sequencing (Plasmidsaurus / Primordium).

To screen the library, a reporter HEK293T cell line carrying a single-copy genomically integrated EF1α-driven destabilized mCitrine (mCitrine-PEST) cassette was generated by AAVS1 HDR and isolated as a monoclonal line by FACS. Library members were transiently transfected using Fugene HD (Promega E2311) into the reporter cell line, each carrying a crRNA targeting the EF1α promoter or a non-targeting (NT) control. Reporter mCitrine fluorescence was monitored by flow cytometry at 48, 72, 96, 120, 144, 168, 264, 312, and 360 hr post-transfection, and two quantitative metrics were derived for each variant: repression potency (median fold-reduction in mCitrine at peak repression), persistence (median fold-reduction at 360 hrs post-transfection), and specificity (On-target repression with EF1α promoter targeting crRNA normalized to Off-target repression with NT crRNA).

### Routine molecular cloning

All plasmids were constructed using Gibson assembly (NEBuilder HiFi DNA Assembly Master Mix, NEB E2621) or Golden Gate cloning (NEBridge Golden Gate Assembly Kit (BsmBI-v2), NEB E1602). Fragments were generated by PCR using Q5 High-Fidelity DNA Polymerase (NEB M0492) or by gene synthesis (IDT gBlocks). Assemblies were transformed into NEB Stable competent E. coli (NEB C3040) for lentiviral and repeat-containing constructs, or into NEB 10-beta (NEB C3019) for standard cloning. Plasmid DNA for transfection was prepared with the QIAprep Spin Miniprep Kit (Qiagen 27104).

### crRNA and poly-crRNA array design

Individual crRNAs were designed primarily using the Broad Institute CRISPick web portal (https://portals.broadinstitute.org/gppx/crispick/public) with the following settings: reference genome matched to the target species (GRCh38 for human; GRCm39 for mouse), Mechanism = CRISPRi, Enzyme = AsCas12a, with all other parameters left at default. The ‘Input’ (target gene) and ‘sgRNA Sequence’ columns from the CRISPick picking results were exported and used as the primary crRNA pool. For each receptor, 3–5 top-ranked crRNAs were selected (full list in [SI Table — crRNA Sequences]).

For genes for which the CRISPick portal failed to return candidate crRNAs, crRNAs were designed manually using the following procedure. First, the gene of interest was located on the UCSC Genome Browser (https://genome.ucsc.edu/) using the species-matched reference genome. The transcription start site (TSS) was identified from the annotated gene track (e.g., for ACVR1C, the TSS corresponds to the right edge of the gene track on the minus strand). Genomic coordinates spanning −200 to +200 bp relative to the TSS were extracted. These coordinates were then used as input to the Benchling CRISPR design tool (Import from: Chromosomal Coordinates; reference genome matched to UCSC). Within Benchling, the entire imported sequence was selected as the ‘Target region’ and the CRISPR guide design tool was run with AsCas12a / CRISPRi-compatible settings (TTTV PAM, 20-nt spacer). Candidate crRNAs were curated manually based on the Benchling on-target score and inter-crRNA spacing across the −200 to +200 bp window, and exported. Manually designed crRNAs were combined with the CRISPick-derived pool to generate the final per-gene crRNA list.

Poly-crRNA arrays were designed as concatenation between individual 20-nt spacers separated by the LbCas12a 20-nt direct repeat (DR; 5′-AATTTCTACTAAGTGTAGAT-3′). For arrays of ≥15 crRNAs, 4-bp synthetic separators (syn-sep; AT-rich sequences inspired by the natural Cas12a DR flanking context, sequence: 5′-AAAT-3′) were inserted on both sides of every crRNA spacer to enhance pre-crRNA processing fidelity (Magnusson et al., 2021). Arrays shorter than 200 bp were synthesized as IDT ultramers; arrays longer than 200 bp were either assembled from multiple IDT ultramers (≤200 bp) with computationally designed compatible sticky ends using Golden Gate cloning, or directly synthesized from Genscript.

The 52-crRNA pan-receptor array used to deplete all twelve expressed TGF-β superfamily receptors (Fig. 3H) targets BMPR1A, BMPR1B, BMPR2, ACVR1, ACVR1B, ACVR1C, ACVR2A, ACVR2B, TGFBR1, TGFBR2, TGFBR3, and ACVRL1. To generate diverse receptor-subset configurations (B_off_, T_off_, A_off_, A_only_) from this parental array, we developed a modular combinatorial assembly strategy based on slot-encoded binary values (Fig. S5B). Each receptor’s crRNA cassette (containing all crRNAs targeting that receptor, flanked by direct repeats) was prepared in advance as a re-usable PCR amplicon. In parallel, a length-matched ‘spacer filler linker’ amplicon containing no functional crRNA was prepared for each slot. Each receptor configuration was then encoded as a binary vector across the twelve receptor slots, with a value of 1 (receptor present in the array, i.e., targeted) directing inclusion of the corresponding crRNA-cassette amplicon and a value of 0 (receptor absent, i.e., un-targeted) directing inclusion of the spacer filler at that slot. Slot-specific Golden Gate overhangs (BsmBI) on flanking amplicons enforced positional ordering, allowing a single combinatorial Golden Gate reaction to assemble the chosen amplicon set into the backbone in the correct order. This modular strategy allowed any of 2^12^ possible receptor configurations to be assembled from the same library of pre-amplified DNA components within one week, without re-synthesis of the array (Fig. S5B).

### AAVS1 safe-harbor HDR engineering of Pathway Sculptor lines

*Pathway Sculptor* cell lines were generated by CRISPR/Cas9-mediated homology-directed repair (HDR) at the human AAVS1 safe-harbor locus. The donor plasmid carries (in this order, 5′ → 3′ between AAVS1 homology arms of ∼800 bp each): a splice-acceptor SA-2A-PuroR cassette enabling promoter-less selection (resistance is expressed only upon correct integration into the AAVS1 first intron), a bidirectional TRE3G ↔ CAG promoter driving in one direction poly-cistronic Tet-On-3G transactivator and poly-crRNA array separated by RNA triplex, and in the opposite direction the stability-enhanced ZIM3-KRAB-dCas12a fusion.

PDG458 (Addgene #100900; Cas9 + AAVS1 sgRNA) and the donor plasmid were co-delivered at 1:1 mass ratio, 500 ng total per well of 24-well plate) by Fugene HD. 48 hr post-delivery, cells were placed under puromycin selection (HEK293T: 1 µg/mL; NMuMG: 2 µg/mL) for 3–5 days until untransfected control wells were fully cleared. The bidirectional architecture and promoter-less SA-2A-PuroR design together yielded >80% integration-correct polyclonal populations, allowing direct use without monoclonal derivation for HEK293T-based experiments.

### Transient chemical transfection

Routine transient transfection of all-in-one constructs into HEK293T and dual-reporter HEK293T cells was performed with Fugene HD (Promega E2311) per manufacturer instructions. Briefly, cells were plated at 40% confluency in 96-well (per-well: 100 ng plasmid, 0.6 µL Fugene HD, 10 µL Opti-MEM) or 24-well (per-well: 500 ng, 3 µL Fugene HD, in 50 µL Opti-MEM) format the day before transfection. Transfection complexes were assembled in Opti-MEM, incubated for 15 min at room temperature, and added drop-wise to cells.

### Library preparation for mRNA-sequencing

mRNA were extracted from 96-well plate using Direct-zol-96 RNA Kits (Zymoresearch Cat# R2055). 50 ng of extracted mRNA from each sample were used as inputs for downstream NGS library preparation.

mRNA-seq library was prepared in 96-well format with a modified 3’Pool-seq protocol (https://bmcgenomics.biomedcentral.com/articles/10.1186/s12864-020-6478-3#Sec9). In brief, reverse transcription reaction was prepared by mixing input RNA with 1 μl Indexed RT Primer (10 μM), 1 μl 10 mM dNTP Mix (New England Biolabs Cat# N0447S),1 μl diluted ERCC Spike-In Mix 1 (0.004 μL stock ERCC per μg RNA, Thermo Fisher Scientific Cat# 4456740), 3.6 ul of 5x RT buffer (Thermo Fisher Scientific Cat# EP0752), 0.5 ul of RNase inhibitor (Thermo Fisher Scientific Cat# EO0381), 1 ul Maxima RT H minus (Thermo Fisher Scientific Cat# EP0752), 2.5 ul 10 uM Template Switching Oligo into a 18 ul reaction. Reverse transcription was carried out in a thermocycler with program described in 3’Pool-seq protocol.

Samples from each row of 96-well plate were pooled (column pooling) by mixing an equal volume of each Reverse Transcription reaction into a new well at a total volume of 20 μl. Residual primers were then degraded with the addition of 1 μl Exonuclease I (New England Biolabs) and incubated at 37 °C for 45 min followed by denaturation at 92 °C for 15 min. Subsequent cDNA amplification, tagmentation, and row pooling was performed following 3’Pool-seq protocol.

Finally, 20 ul of pooled NGS library was subject to Gel-based size selection using E-Gel EX Agarose Gel (Thermo Fisher Scientific Cat# G401001) to enrich for fragments with size range between 200-1000 bp and eluted in 15 ul.

Eluted pooled NGS library was examined in an Agilent TapeStation 4200 (Agilent Technologies) to determine average fragment sizes. Library concentration was quantified in a Qubit 3.0 Fluorometer (Life Technologies). NGS library molarity was then calculated using 660 g/mol per base-pair as a molecular weight. NGS library was diluted to 2 nM, denatured in 0.2N NaOH, and loaded onto Element AVITI sequencer following Element Biosciences Cloudbreak Sequencing user guide.

### Flow cytometry

Cells for assessment were trypsinized with 40 μL of 0.05% trypsin-EDTA (Gibco) for 1 minute at room temperature, and subsequently resuspended in 100 μL of Hanks’ Balanced Salt Solution (HBSS) containing 2.5mg/ml bovine serum albumin (BSA), 1mM ethylenediaminetetraacetic acid (EDTA), and/or 4 units/ml DNase I (NEB). Cells were then filtered through a 40 μm cell strainer (Falcon™) or a 96-well plate cell strainer (Millipore) and analyzed by flow cytometry (CytoFLEX, Beckman Coulter or ZE5, Bio-Rad).

### RT-qPCR

Total RNA was harvested from cell lysate using the RNeasy mini kit (QIAGEN) and cDNA was generated from 1μg of RNA using the iScript cDNA synthesis kit (BioRad) following the manufacturer’s instructions. Primers and probes for specific genes were purchased from IDT. Reactions were performed using 1:40 dilution of the cDNA synthesis product with either IQ SYBR Green Supermix or SsoAdvanced Universal probes Supermix (BioRad). Cycling was carried out on a BioRad CFX96 thermocycler using an initial denaturing incubation of 95° for 3 min followed by 39 cycles of (95°C for 15 s, followed by 60°C for 30 s). Each condition was assessed with three biological repeats and each reaction was run at least in triplicate.

### Dual BRE/TRE reporter cell line construction

To monitor BMP- and TGF-β-branch signaling in the same cell, a dual reporter cassette was constructed encoding (i) a 12×BRE element (BMP-responsive element from the Id1 promoter; Korchynskyi & ten Dijke, 2002) driving mCitrine-PEST from the minimal Major Late Promoter (MLP) promoter, and (ii) a 12×TRE element (CAGA-box from the PAI-1 promoter; Dennler et al., 1998) driving mCherry-PEST from a minimal MLP promoter, flanked by the insulator cHS4. This cassette was inserted between PiggyBac inverted repeats and integrated into HEK293 cells as described under “Cell line engineering” section. Following selection with hygromycin and puromycin, monoclones were isolated by FACS and assessed for dynamic range (fold-change in reporter signal upon stimulation with select BMP ligands for BRE or select TGF-β ligands for the TRE at 24 hr). The selected dual-reporter HEK293T clone was validated by demonstrating dose-dependent activation of each reporter by its cognate ligand panel with high signal-to-noise (Fig. 4B).

An analogous dual-reporter NMuMG cell line (NAMRU mouse mammary gland epithelial; ATCC CRL-1636; cultured in DMEM + 10% FBS + 10 µg/mL insulin + 1× Pen/Strep, passaged every 2–3 days) was engineered by the same strategy. Monoclones were isolated and validated against the corresponding mouse ligand panel.

### Cell surface staining

Cells for assessment were dissociated with accutase (Stemcell Technologies Cat# 07922) for 5min at RT and neutralized with 200 μL of Hanks’ Balanced Salt Solution(HBSS) containing 2.5mg/ml bovine serum albumin (BSA), 1mM ethylenediaminetetraacetic acid (EDTA), and/or 4 units/ml DNase I (NEB). Dissociated cells were filtered with a 40 um cell strainer and spun down at 300g for 5 min in V-bottom 96-well plate. Cells were then resuspended with flow buffer containing corresponding 1:500 diluted FP-conjugated antibodies targeting CD markers for 1 hr at RT on a belly dancer. After staining, cells were spun down and washed with flow buffer once before loading onto cytoflex machine for fluorescence measurements.

### Cell line engineering

Safe-harbor locus targeting: PDG458 (Addgene Plasmid #100900) with sgRNA targeting rosa/AAVS1 locus and donor plasmids with corresponding homology arms flanking sculptor and rtta were co-delivered into cells with either chemical transfection or LNP. Cells were subject to antibiotic selection 48 hrs post-delivery and were continuously cultured in selection media for 3-7 days (cell bottlenecking), depending on the administered antibiotics. A positive control condition where either the Cas9 plasmid or donor plasmid were omitted in the delivery were used to indicate the completion of cell bottlenecking. Specifically for mESC, monoclones with appropriate expression level were derived by sorting single cells with flow cytometer and expanded for 1 week. For all other cell types, polyclonal population derived from antibiotic-based bottlenecking were directly used for experiments. We note that, as opposed to a PB-based random integration strategy, targeted strategy yields a relatively homogeneous population and therefore, for most of the cases, monoclonal derivation can be omitted.

### TGF-β superfamily ligand stimulation

Recombinant ligands sources, catalog numbers, and reconstitution conditions: TGF-β1 (R&D Systems 7666-MB-005), TGF-β2 (R&D 7346-82-005), TGF-β3 (R&D 243-B3), Activin A (R&D 338-AC-010), Activin B (R&D 8260-AB-010), GDF11 (R&D 1958-GD-010), Nodal (R&D 3218-ND), and the BMP ligands BMP4 (R&D 5020-BP-010), BMP5 (R&D 6176-BM-020), BMP9 (R&D 5566-BP-010), BMP10 (R&D 6038-BP-025), and GDF5 (R&D 853-G5-050). All ligands were reconstituted in 4 mM HCl / 0.1% BSA per manufacturer instructions and stored at −80 °C in single-use aliquots.

Cells were plated at 40% confluency in 96-well plates and cultured for 12 hr. Medium was replaced with fresh medium containing the indicated ligand at the indicated final concentration. For dose-response curves (Fig. 4B, 5D), ligands were applied across [6, 3-fold serial dilutions spanning 8 - 2000 ng/mL or 0.02 - 5 ng/mL (TGF-β1, TGF-β2)]. For all other ligand experiments, ligands were applied at saturating concentrations (typically [200-250 ng/mL or 5 ng/mL (TGF-β1, TGF-β2)]). Cells were harvested for flow cytometry 24 hr after ligand addition unless otherwise noted. For combined ligand experiments (Fig. 6B, ActB+BMP9 antagonism), each ligand was applied alone or together at the same per-ligand saturating concentration.

### mRNA preparation and receptor reconstitution

Mouse Acvr1 and Acvr2a mRNAs were generated by in vitro transcription (IVT). Coding sequences (mouse Acvr1, NM_007394; mouse Acvr2a, NM_007396) were PCR amplified using a 5’ primer with a T7 promoter addition and The HiScribe T7 ARCA mRNA Kit with tailing (NEB E2060S) RNA was purified using RNeasy Mini cleanup (QIAGEN), quantified with an 8 channel Nanodrop (Thermo) and analyzed for integrity using 2% agarose gel with SYBR Gold (Thermo).

For receptor reconstitution (Fig. 4C–E), Pathway Sculptor cells with all BMP receptors knocked down (induced with 1 µg/mL Dox for 72 hr) were plated in 96-well format. The indicated dose of Acvr1 mRNA + Acvr2a mRNA (0, 7, 14, 28, 56, 112 ng each per well) was complexed with Lipofectamine MessengerMAX (Thermo Fisher LMRNA001) in Opti-MEM at a 3 µL MessengerMAX : 1 µg mRNA ratio per manufacturer instructions, and applied to cells. After 24 hr, medium was replaced with ligand-containing medium, and cells were harvested for flow cytometry 24 hr post-stimulation. Reporter response was normalized to no-receptor (0 ng) control.

### Primary bone marrow cell electroporation

Primary bone marrow cells expressing CD19, CD34, and CD326 were obtained from the crushed bones of mice, and primary human cells expressing CD55 and CD321 were obtained from mobilized peripheral blood from anonymized donors. Both cell populations were isolated via FACS sorting with a FACS-Aria II (BD Biosciences). An all-in-one plasmid DNA construct expressing a poly crRNA array targeting the respective CD markers and Sculptor were electroporated into the cells using a Neon Electroporation System (Thermo Fisher Scientific) with Buffer T, on the following parameters: 1700 V, 20 ms, 1 pulse. Following electroporation, the cells are cultured in RPMI 1640 (Gibco) at 37 °C, 5% CO_2_ for 96 hours. The median fluorescence intensity (MFI) of each CD marker is quantified via FACS analysis with the Attune Nxt (Invitrogen).

### Quantifications and statistical analyses

Flow cytometry data analyses: Flow cytometry data was analyzed in MATLAB using a custom software (EasyFlow: <u>GitHub - AntebiLab/easyflow: Matlab Based Flow Cytometry Analysis Tool</u>). Events collected from flow cytometry experiments were first gated based on forward vs. side scatter to select for cells, followed by gating based on forward scatter area vs forward height, to select for single cells.

mRNA-seq analyses: Read de-multiplexing was performed with Bases2Fastq, a standard software package used by Element Bioscience system. Reads were aligned to a custom reference genome GRCh38.103 using STAR (2.7.8a) (https://www.sciencedirect.com/science/article/pii/S009286742300689X?via%3Dihub#bib78) with the ENCODE standard options except “--outFilterScoreMinOverLread 0.3 --outFilterMatchNminOverLread 0.3 --outFilterMismatchNmax 20 –outFilterMismatchNoverLmax 0.3 --alignSJoverhangMin 5 --alignSJDBoverhangMin 3”. Uniquely mapped reads that overlap with genes were counted using HTSeq-count (0.13.5) (https://www.sciencedirect.com/science/article/pii/S009286742300689X?via%3Dihub#bib79) with default settings except “-m intersection-strict”. Since 3’Pool-seq sequences only the 3′-end of mRNA transcripts, no gene length normalization was applied to read counts when calculating Transcripts Per Million (TPM) values. Differential gene expression analysis was carried out using the DESeq2 package in R (https://genomebiology.biomedcentral.com/articles/10.1186/s13059-014-0550-8).

### Dose-response curve fitting

Reporter fold-change vs. ligand concentration data (Figs. 4B, 4E, 5D, S6) were fit to one of two parametric models, with model selection by corrected Akaike information criterion (AICc):

i. Hill (monotonic) model: y(c) = y_0_ + (y_max_ − y_0_) · c^n^ / (K^n^ + c^n^), with free parameters y_0_ (basal), y_max_ (saturation), K (EC_50_), n (Hill coefficient).
ii. Biphasic model (sum of an activating and an inhibitory Hill term): y(c) = y_0_ + (y_max_ − y_0_) · c^n1^ / (K_1_^n1^ + c^n1^) · K_2_^n2^ / (K_2_^n2^ + c^n2^).

Both models were fit by non-linear least squares using scipy.optimize.curve_fit (SciPy v1.11) on log_10_-transformed concentrations, with parameter bounds n ∈ (0.5, 5), K ∈ (10^−3^, 10^3^) ng/mL, y_max_ > y_0_. Initial parameter values were set heuristically from the data extrema. Each ligand–receptor configuration was fit independently across three biological replicates (mean ± s.d. plotted; lines show the fit to the mean). The biphasic model was selected only when ΔAICc > 4 favored it over the Hill model and when the fitted K2 was within the experimental concentration range. Code is available upon request.

### Construction of 2-D ligand vectors

To represent each ligand’s signaling effect as a 2-D vector, BRE-YFP and TRE-RFP responses were log-transformed and rescaled per receptor configuration as follows. For each receptor configuration *p* (wild-type, B_off_, T_off_, A_off_, A_only_), each Dox condition *d*, and each ligand *i*, the per-replicate fold-change relative to the unstimulated control at saturating ligand concentration was computed for both reporters, *F*_BRE_(*p,i,d*) and *F*_TRE_(*p,i,d*). Fold-changes were then log_2_-transformed after addition of a small pseudocount (ε = 0.01) to stabilize values near zero:

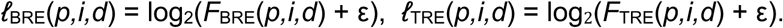

Per-replicate uncertainties on the linear fold-change, σ_F_, were propagated to the log scale as σ_ℓ_ = σ_F_ / [(*F* + ε) · ln 2].

To place BRE and TRE on a comparable scale, each axis was rescaled by its standard deviation computed *within* the receptor configuration, pooling across all ligands and Dox conditions for that perturbation:

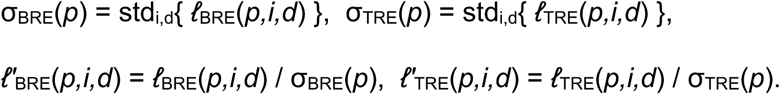

Here the prime denotes the per-perturbation σ-rescaled log-fold-change (corresponding to BRE_norm / TRE_norm in the analysis code). Importantly, scaling is performed independently per receptor configuration (rather than by a wild-type-only maximum). This per-perturbation rescaling expresses each axis in units of its own variability and allows direct comparison of ligand identity within each receptor configuration. Propagated uncertainties on the rescaled axes are σ_ℓ′_ = σ_ℓ_ / σ_axis_(*p*). No mean-centering is applied, so the origin retains its biological meaning of “no log-fold change above background.”

The resulting 2-D ligand vector used for all downstream analyses is

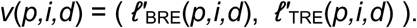

### Cosine similarity computation

Cosine similarities between ligand pairs within a (*p,d*) condition, and between Dox+ and Dox− conditions of the same ligand, were computed directly on these vectors using sklearn.metrics.pairwise.cosine_similarity. Family-averaged cosine similarities were obtained by averaging individual ligand-pair (or Dox-pair) cosine similarities within each ligand family or perturbation × Dox stratum.

### Interaction decomposition (additivity analysis)

To assess whether the BMP and TGF-β receptor families contribute additively or synergistically to a given reporter response (Figs. 5H, 5I, S6B), each ligand’s saturating-dose reporter fold-change was decomposed into baseline, single-branch, and interaction terms. Let R_WT, R_T_off_, R_B_off_, and R_KD denote linear-scale fold-change in wild-type, T_off_ (BMP-receptors-only), B_off_ (TGF-β-receptors-only), and full-knockdown cells (assumed to be no response), respectively. Each is normalized to its own unstimulated control. The decomposition is:

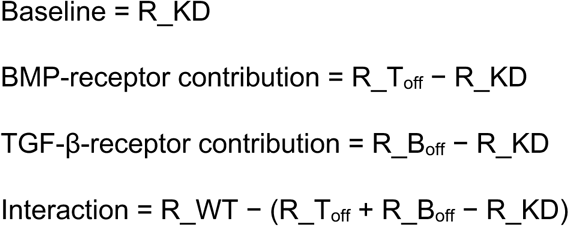

A positive interaction term indicates super-additive (synergistic) cooperation; a negative term indicates sub-additive (antagonistic) interaction.

For the ligand antagonism analysis (Fig. 6B), where two ligands (A and B) were applied alone or in combination at saturating concentrations, the interaction term was computed in log_2_ space, as a fractional (rather than absolute) deviation of the observed combination response from a multiplicative-independence null. Let R_A, R_B, R_AB, and R_NL denote the per-replicate fold-change of the reporter relative to its unstimulated baseline for ligand A alone, ligand B alone, the A+B combination, and the no-ligand control, respectively. Under multiplicative independence — i.e., the assumption that the two ligands’ effects compound proportionally — the expected log_2_ fold-change of the combination is the additive sum of the individual log_2_ fold-changes:

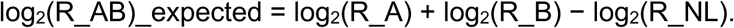

The log-space interaction term is the deviation of the observed log_2_ combination response from this expectation:

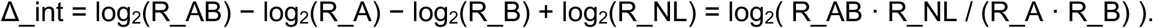

By construction, Δ_int = 0 corresponds to multiplicative independence (no interaction), Δ_int > 0 indicates super-additive (synergistic) cooperation, and Δ_int < 0 indicates sub-additive (antagonistic) interaction. We use this log-space (fractional) formulation rather than a linear “observed minus (A + B − NL)” or “observed minus (A · B / NL)” deviation because a log-fold-change deviation measures the relative shift away from the expected response, which is invariant to the absolute magnitude of the expected response and therefore comparable across receptor configurations and ligand pairs whose dynamic ranges differ. This matters when comparing perturbations (e.g., Dox-induced receptor knockdowns) that change the baseline expected combination response by large factors: the same absolute deviation can correspond to very different fractional deviations, and only the latter reflects the biological strength of the interaction.

