## Supplemental Figures for "Programmable pathway profiles reveal signaling principles of TGF-β superfamily receptors"

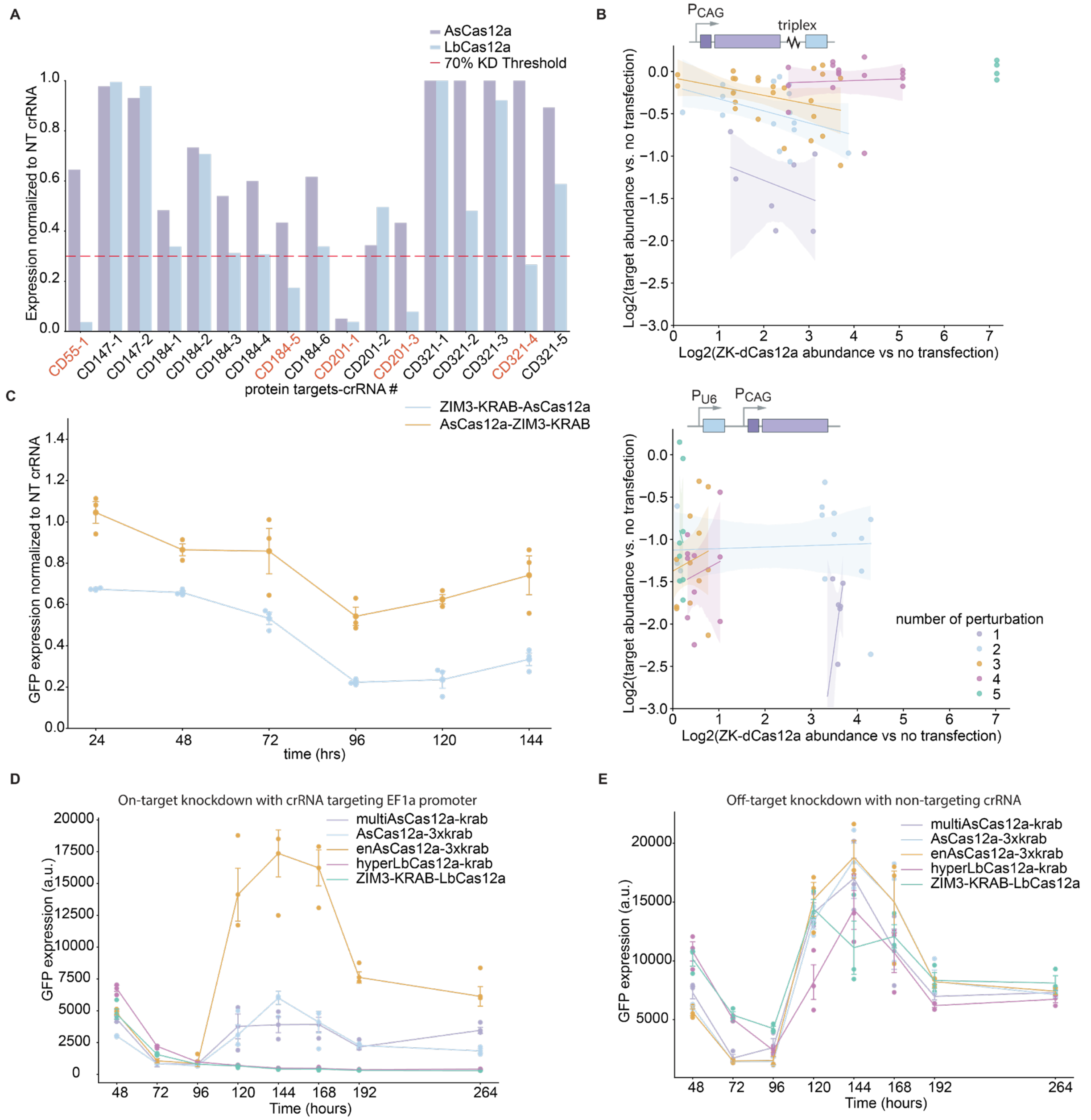


**Fig. S1**. **Design space of an enhanced dCas12a-effector.** Optimized parameters include **(A)** Comparison of LbdCas12a- vs. AsdCas12a-KRAB repression efficiencies of a panel of CD surface proteins. **(B)** crRNA abundance vs. knockdown correlation. Top: A construct expressing a ZK-dCas12a and a poly-crRNA array in a coupled manner, where poly-crRNA expression level is constrained. Bottom: a construct expressing a ZK-dCas12a and a poly-crRNA array in a de-coupled manner, where poly-crRNA expression is unconstrained. (**C**) Comparison of N- vs. C-terminal KRAB fusion repression efficiencies of mCitrine reporter. (**D, E**) On-target/off-target KD time courses across five dCas12a-effector variants.

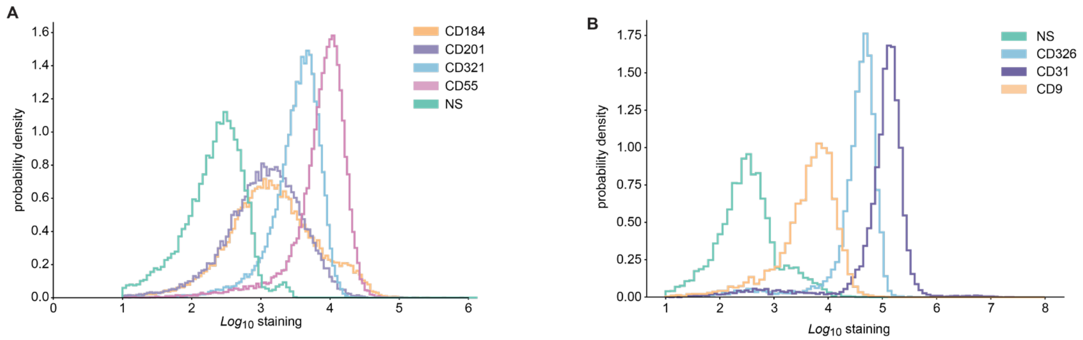


**Fig. S2**. **Basal CD surface marker expression distributions.** (**A**) Flow cytometric measurement of antibody staining of CD184, CD201, CD321, CD55 in HEK293T. NS: no-staining control. (**B**) Flow cytometric measurement of antibody staining of CD326, CD9, CD31 in mESC. NS: no-staining control.

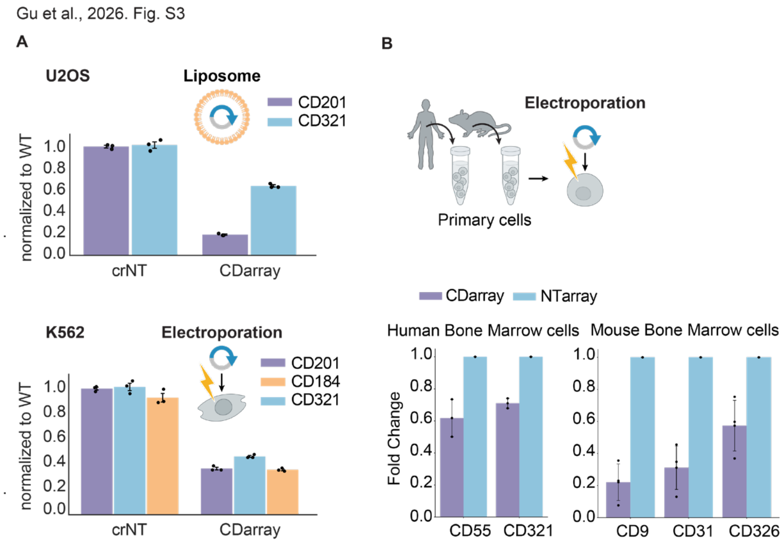


**Fig. S3.** ***Pathway Sculptor* is robust across diverse cell types and delivery approaches. (A)** Flow cytometric measurement of the expression levels of surface proteins, as indicated in **Fig. 2A**. Dot denotes median staining intensity normalized to WT from independent biological replicates. Bar heights and error bars denote the mean ± s.e.m. across $n$ = 3 independent biological replicates. **(B)** Flow cytometric measurement of surface protein expression as in **Fig. 2B** (human) and **Fig. 2G** (mouse), upon electroporation of an all-in-one *Pathway Sculptor* construct encoding poly-crRNA arrays targeting human or mouse CDs into bone marrow cells from human and mouse donors, respectively.

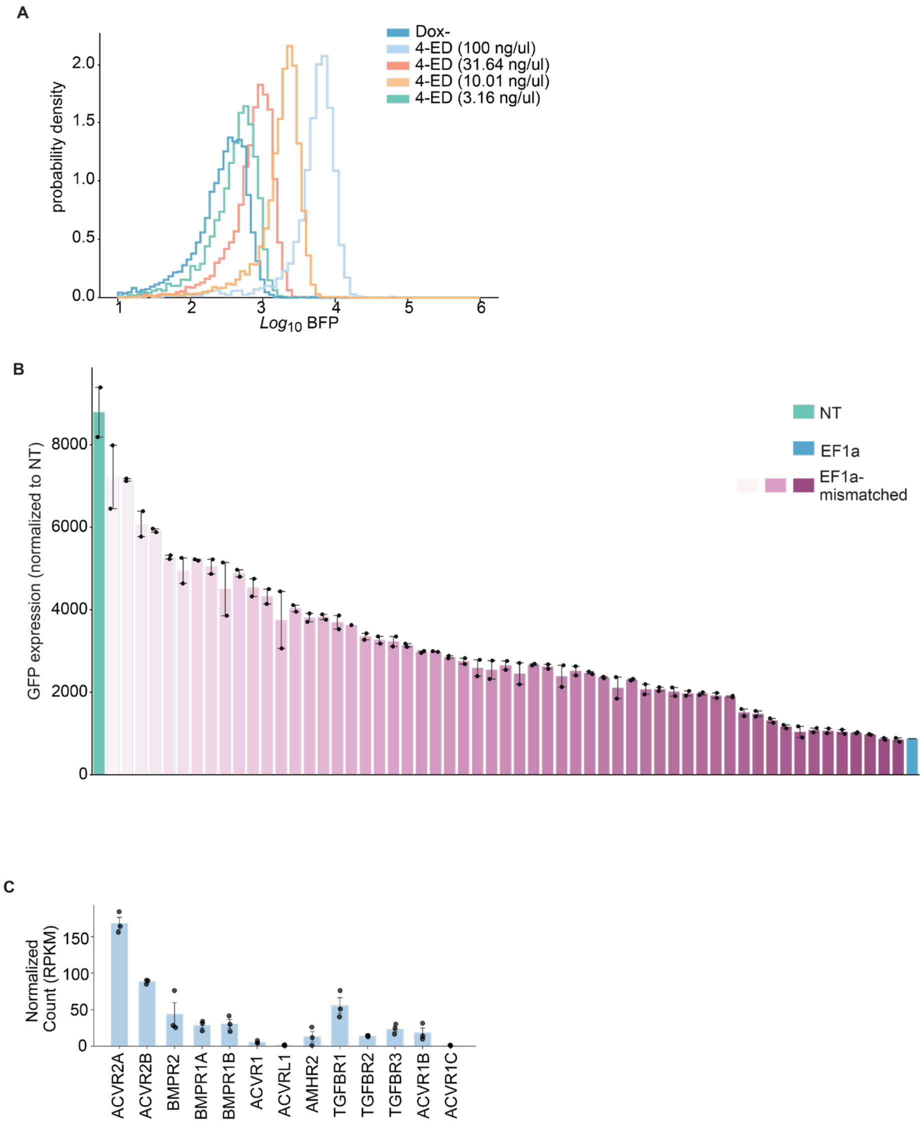


**Fig. S4**. **Tunability of *Pathway Sculptor*-mediated knockdown.** (**A**) 4-epidoxycycline dose-response histogram in stable HEK293 cells. (**B**) Tunable modulation of EF1α-citrine expression by mismatched crRNA panel demonstrating the ≥2-mismatch knockdown. (**C**) Transcription abundance of all twelve TGF-β superfamily receptors in HEK293T cells.

Gu et al., 2026. Fig. S5

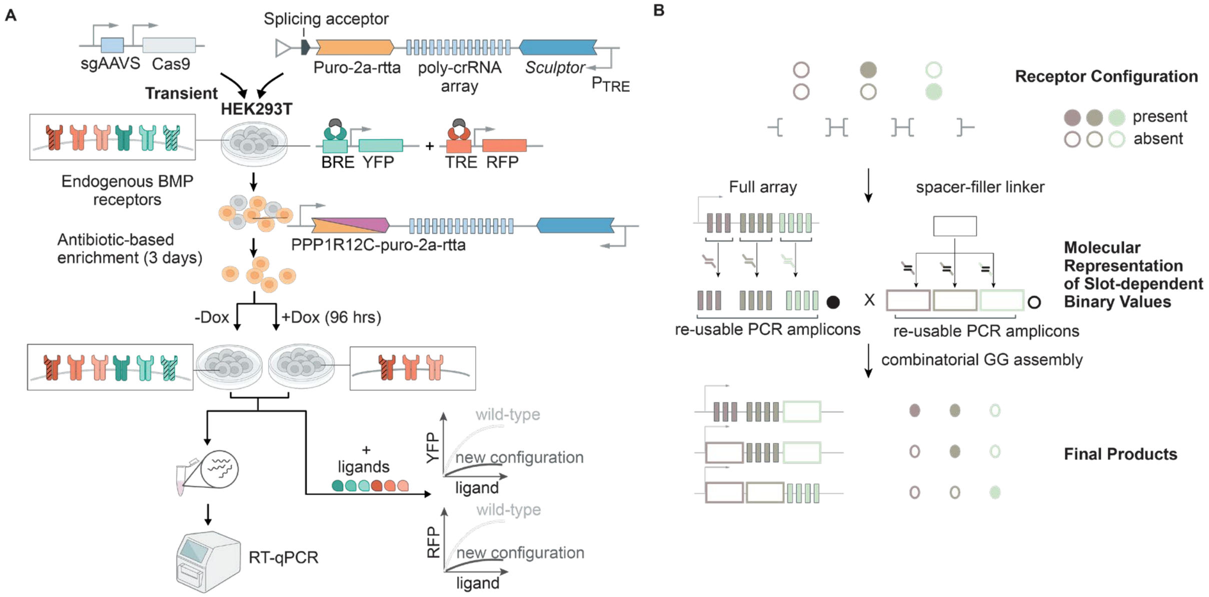


**Fig. S5**. **Stable cell line engineering and combinatorial poly-crRNA array assembly strategy. (A)** Schematics of engineering of cell lines carrying BRE/TRE dual-reporter and AAVS-targeted *Pathway Sculptor.* (**B**) combinatorial assembly strategy for receptor re-configurations.

Gu et al., 2026. Fig. S6

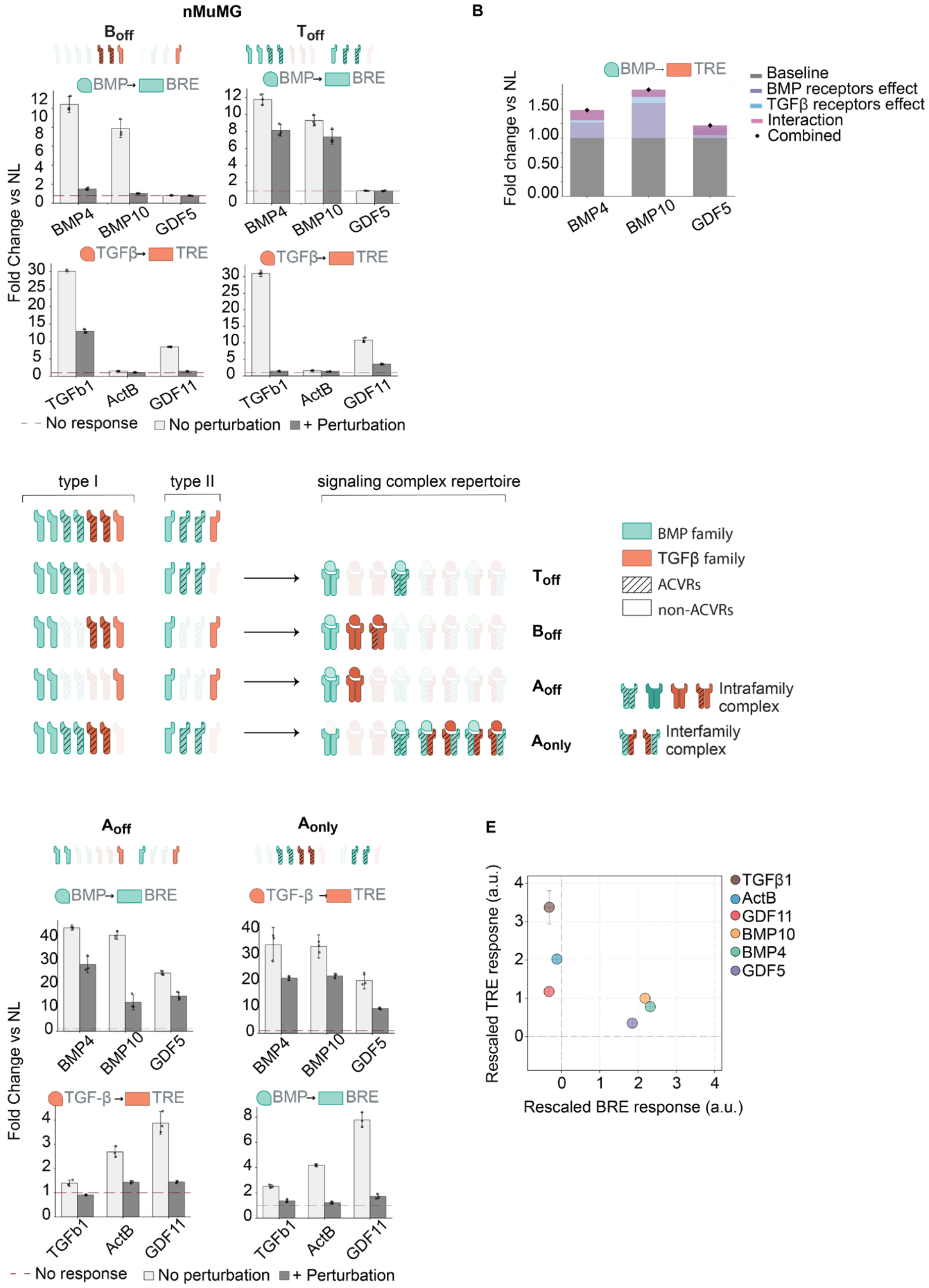


**Fig. S6**. **Extended BMP/TGF-β crosstalk in HEK293 and nMuMG cells**. (**A**) nMuMG ligand responses in $B_{off}$ and $T_{off}$ cell lines. (**B**) receptor contribution decomposition. (**C**) signaling complex repertoire. (**D**) $A_{off}$/$A_{only}$ responses and (**E**) Ligand identity scatter plot.

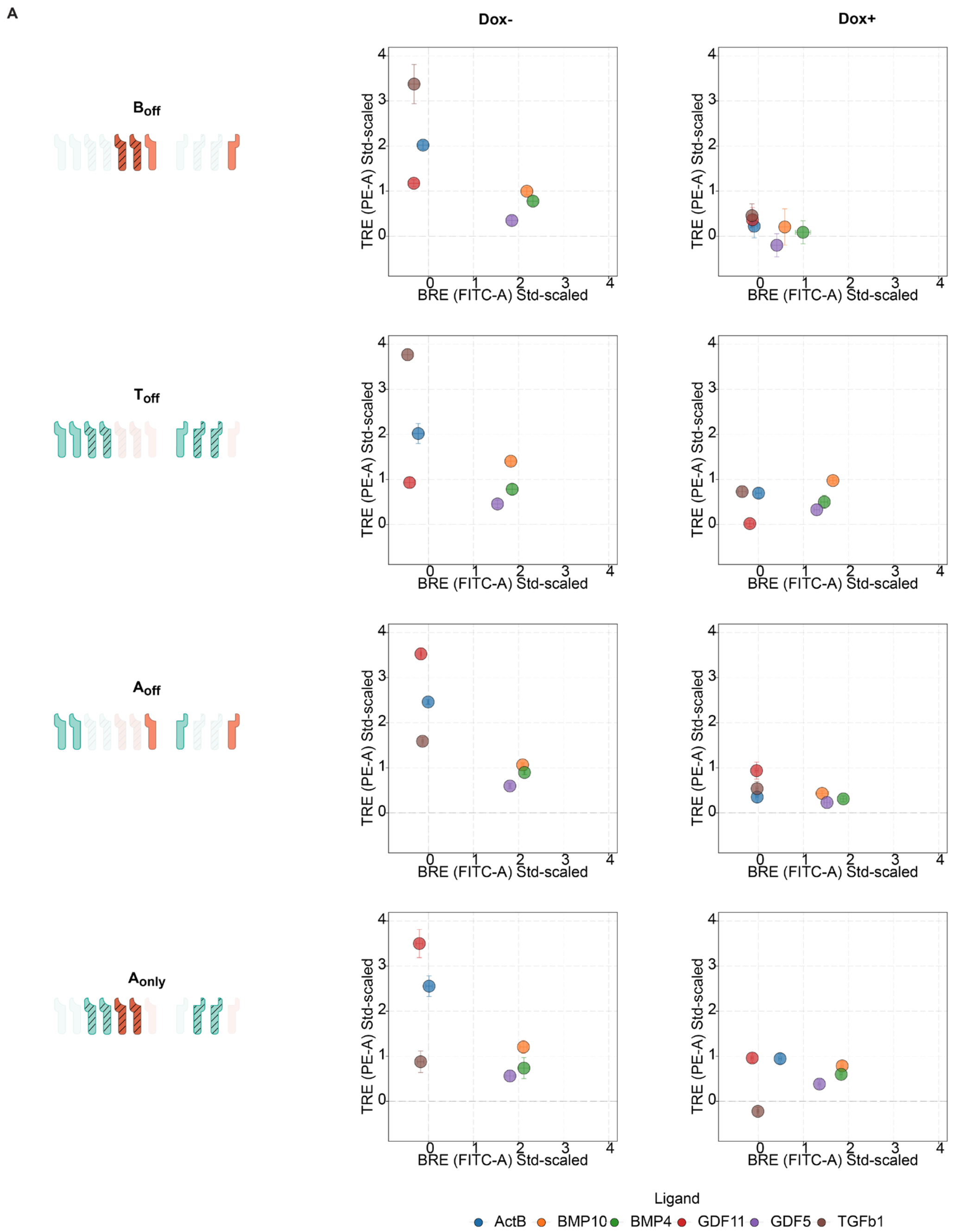


**Fig. S7**. **Ligand response vectors across reprogrammed receptor profiles.** (**A**) 2-dimensional rescaled-TRE/BRE responses for six tested ligands in Dox−/Dox+ conditions across all four receptor configurations.
